# Mechanism and inhibition of SARS-CoV-2 PLpro

**DOI:** 10.1101/2020.06.18.160614

**Authors:** Theresa Klemm, Gregor Ebert, Dale J. Calleja, Cody C. Allison, Lachlan W. Richardson, Jonathan P. Bernardini, Bernadine G. C. Lu, Nathan W. Kuchel, Christoph Grohmann, Yuri Shibata, Zhong Yan Gan, James P. Cooney, Marcel Doerflinger, Amanda E. Au, Timothy R. Blackmore, Paul P. Geurink, Huib Ovaa, Janet Newman, Alan Riboldi-Tunnicliffe, Peter E. Czabotar, Jeffrey P. Mitchell, Rebecca Feltham, Bernhard C. Lechtenberg, Kym N. Lowes, Grant Dewson, Marc Pellegrini, Guillaume Lessene, David Komander

**Affiliations:** The Walter and Eliza Hall Institute of Medical Research and Department of Medical Biology, University of Melbourne, 1G Royal Parade, Melbourne, VIC 3052, Australia, Melbourne; Department of Biochemistry and Molecular Biology, Michael Smith Laboratories University of British Columbia, Vancouver, Canada; Oncode Institute and Department of Chemical Immunology, Leiden University Medical Centre, Einthovenweg 20, 2333 ZC, Leiden, The Netherlands; Commonwealth Scientific and Industrial Research Organisation (CSIRO), Biomedical Program, Parkville, VIC 3052, Australia; Australian Synchrotron, ANSTO, 800 Blackburn Road, Clayton, VIC 3168, Australia; Pharmacology and Therapeutics Department, University of Melbourne, Melbourne VIC 3020, Australia

## Abstract

Coronaviruses, including SARS-CoV-2, encode multifunctional proteases that are essential for viral replication and evasion of host innate immune mechanisms. The papain-like protease PLpro cleaves the viral polyprotein, and reverses inflammatory ubiquitin and anti-viral ubiquitin-like ISG15 protein modifications^1,2^. Drugs that target SARS-CoV-2 PLpro (hereafter, SARS2 PLpro) may hence be effective as treatments or prophylaxis for COVID-19, reducing viral load and reinstating innate immune responses^3^. We here characterise SARS2 PLpro in molecular and biochemical detail. SARS2 PLpro cleaves Lys48-linked polyubiquitin and ISG15 modifications with high activity. Structures of PLpro bound to ubiquitin and ISG15 reveal that the S1 ubiquitin binding site is responsible for high ISG15 activity, while the S2 binding site provides Lys48 chain specificity and cleavage efficiency. We further exploit two strategies to target PLpro. A repurposing approach, screening 3727 unique approved drugs and clinical compounds against SARS2 PLpro, identified no compounds that inhibited PLpro consistently or that could be validated in counterscreens. More promisingly, non-covalent small molecule SARS PLpro inhibitors were able to inhibit SARS2 PLpro with high potency and excellent antiviral activity in SARS-CoV-2 infection models.

---

The COVID-19 pandemic unfolding globally in the first half of 2020 is caused by the novel Coronavirus SARS-CoV-2, and has highlighted, amongst many things, the general lack of antiviral small molecule drugs to fight a global coronavirus pandemic. Proteolytic enzymes are critical for viruses expressing their protein machinery as a polyprotein that requires cleavage into functional units. As a result, viruses with blocked protease activity do not replicate efficiently in cells; this concept extends to coronaviruses ^4^. Drugging the proteases in SARS-CoV-2 is therefore a current focus of concerted global academic and pharma efforts ^3^.

SARS-CoV-2 encodes two proteases, the papain-like protease (PLpro, encoded within non-structural protein (nsp) 3), and 3-chymotrypsin-like ‘main’ protease (3CLpro or Mpro, encoded within nsp5). PLpro generates nsp1, nsp2, and nsp3 (**Figure 1a**) and 3CLpro generates the remaining 13 non-structural proteins. After their generation, the nsps assemble the viral replicase complex on host membranes, initiating replication and transcription of the viral genome^1,5^.

**Figure 1.**
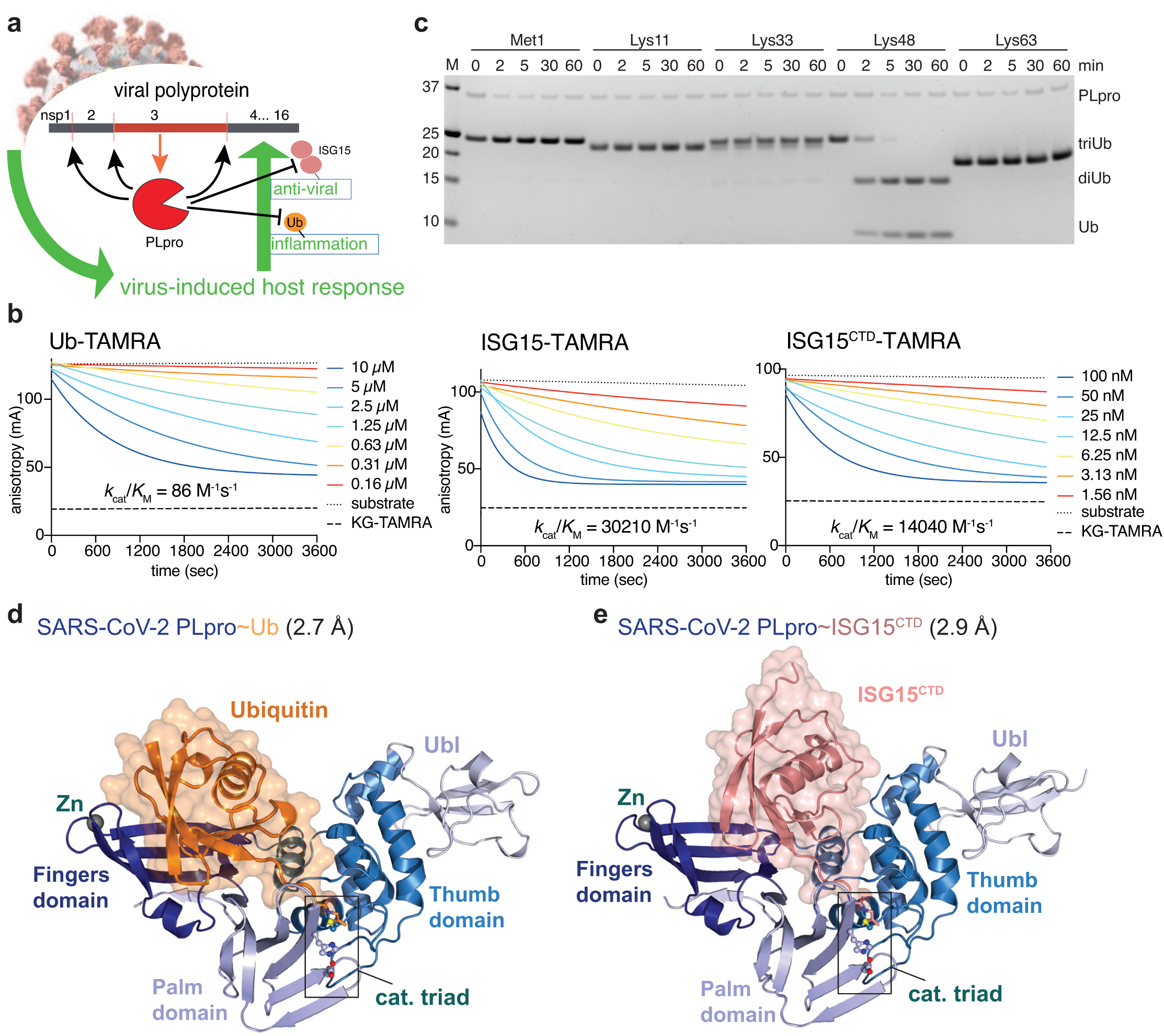
Biophysical and structural characterisation of PLpro activity. **a**, Cartoon of Coronavirus PLpro activities. PLpro is encoded as one of various domains of the 1900 amino-acid non-structural protein nsp3, and is thought to have three functions: (i) cleaving the viral polyprotein to generate mature nsp1, nsp2 and nsp3; (ii) hydrolysing ubiquitin chains important for inflammatory responses, and (iii) removing interferon-stimulated gene 15 (ISG15) modifications from proteins, reversing antiviral responses. **b**, Fluorescence polarisation (FP) assay at indicated PLpro concentrations to derive catalytic efficiency. Experiments were performed in technical triplicate and n=5 (Ub-TAMRA), n= 6 (ISG15-TAMRA), n=4 (ISG15^CTD^-TAMRA) biological replicates with one biological replicate shown as representative. Catalytic efficiencies were calculated as described in **Methods**. Also see **Extended Data Fig. 2. c**, Time course analysis of triubiquitin (2 µM) hydrolysis using 250 nM SARS2 PLpro, resolved on a Coomassie stained SDS-PAGE gel. Linkage specific cleavage of Lys48-linked triubiquitin to di- and monoubiquitin resembles SARS PLpro activity^17,20^. See **Supplementary Figure 1** for uncropped gels, and **Extended Data Fig. 2b-d** for gel-based cleavage quantification. **d**, Crystal structure at 2.7 Å resolution of SARS2 PLpro with subdomains coloured in shades of blue, bound to ubiquitin propargylamine (Ub-PA, orange). Catalytic triad residues are shown in ball- and-stick representation, and a Zn ion is indicated as a grey sphere. Also see **Extended Data Fig. 3** and **Extended Data Table 1. e**, Crystal structure at 2.9 Å resolution of SARS2 PLpro (blue) bound to ISG15 C-terminal domain propargylamide (ISG^CTD^-PA, salmon). Also see **Extended Data Fig. 3** and **Extended Data Table 1**.

Viral proteases can have additional functions, and can for example act to inhibit host innate immune responses that are mounted initially as an inflammatory response, and subsequently as an interferon response. The interferon system generates an antiviral state in host cells through transcriptional upregulation of more than 300 interferon-stimulated genes (ISGs), to efficiently detect and respond to viral threats^6^. Dysregulated inflammatory responses are a hallmark of COVID-19, and substantial morbidity and mortality is associated with overzealous immune responses (a ‘cytokine storm’), causing collateral damage ^7^.

A common mechanism by which viral proteases regulate innate immune pathways is through antagonising ubiquitin and ubiquitin-like modifications (**Figure 1a**) ^8,9^. Protein ubiquitination is complex due to the occurrence of many ubiquitin chain architectures that encode non-degradative and degradative functions^10,11^. Inflammatory signalling pathways rely on distinct ubiquitin signals that are regulated by intricate mechanisms in human cells^12^. ISG15 is a ubiquitin-like (Ubl) modification induced upon viral infection^13^. ISG15 itself comprises two Ubl folds that are fused, structurally resembling diubiquitin^14^. Only few cellular enzymes remove ISG15, enabling this modification to act as a virus-induced danger signal. Importantly, coronaviral PLpro enzymes efficiently remove ISG15 and ubiquitin modifications, dampening inflammation and anti-viral signalling (**Figure 1a**) ^5,15-19^. A large body of work by many laboratories has illuminated SARS and MERS PLpro mechanisms in some detail^1,20^, revealing two binding sites for ubiquitin/Ubl-folds (termed the S1 and S2 ubiquitin binding sites, see ^21^ for nomenclature). We here extend these studies to SARS2 PLpro.

## Biochemical characterisation of SARS2 PLpro activity

SARS2 PLpro is 83% identical to SARS PLpro (**Extended Data Fig. 1**) and is also expected to target ubiquitin and ISG15. In a fluorescence polarisation based quantitative assay^22^, SARS2 PLpro hydrolysed an ISG15-TAMRA fluorescent substrate 350-fold more efficiently as compared to ubiquitin-TAMRA (**Figure 1b, Extended Data Fig. 2a**). Strikingly, a substrate comprising only the C-terminal ubiquitin-like fold of ISG15, (ISG15^CTD^ -TAMRA) was still cleaved with 160-fold higher efficiency compared to ubiquitin (**Figure 1b, Extended Data Fig. 2a**). ISG15 versus ubiquitin preference is hence mediated by recognition of one Ubl-fold, which would bind in the S1 ubiquitin/Ubl binding site of PLpro.

An S2 ubiquitin binding site enables SARS PLpro to preferentially cleave Lys48-linked polyubiquitin^17,20^. A putative S2 site is conserved in SARS2 PLpro (**Extended Data Fig. 1**). Gel-based analysis of different triubiquitin chains reveals very high activity and striking Lys48-polyubiquitin specificity in SARS2 PLpro (**Figure 1c**). In fact, Lys48-triubiquitin is hydrolysed with similar activity compared to cleavage of proISG15 to mature ISG15 in gel-based assays (**Extended Data Fig. 2b-d**). Hence, while the S1 site of PLpro prefers ISG15 modifications, the S2 site reinstates efficient cleavage of Lys48-polyubiquitin to create balanced PLpro activity towards modifiers, in an elegant mechanism of attaining polyubiquitin targeting specificity^21^.

## Structural analysis of SARS2 PLpro ubiquitin and ISG15 complexes

Differential cleavage of ubiquitin and ISG15^CTD^ substrates (**Figure 1b**) indicated that the S1 PLpro interacts with both modifiers distinctly. SARS2 PLpro crystal structures covalently bound to ubiquitin-propargylamide (Ub-PA) at 2.7 Å and to ISG15^CTD^-PA at 2.9 Å resolution, enable direct comparison (**Figure 1d**,**e, Extended Data Table 1, Extended Data Fig. 3**). In concordance with previous structures of SARS and MERS PLpro (**Extended Data Fig. 4**), and conceptually resembling human ubiquitin specific protease (USP) enzymes^21^, PLpro binds ubiquitin in an ‘open hand’ architecture, in which the ubiquitin sits on the ‘Palm’ subdomain, and is held in place by the zinc-binding ‘Fingers’ subdomain, such that the ubiquitin C-terminus, the site of hydrolysis, reaches into the catalytic centre (**Figure 1d, Extended Data Fig. 3a**). The structure of SARS2 PLpro, and the position and orientation of the bound ubiquitin molecule, are highly similar to SARS PLpro∼Ub (pdb 4m0w, ref. ^23,24^, RMSD of 0.54 Å for PLpro, see **Extended Data Fig. 4**).

While ISG15^CTD^ sits similarly on the Palm subdomain, it interacts with the Thumb rather than the Fingers of SARS2 PLpro (**Figure 2a, Extended Data Fig. 5a**). The resulting ∼40° rotation of the Ubl-fold α-helix compared to ubiquitin, leads to shifts of up to 15 Å for structurally identical residues in the Ubl fold (**Figure 2a, Extended Data Fig. 5a,b**). Key interaction sites mediating ISG15^CTD^ - PLpro contacts are centred around ISG15 Trp123 and Pro130/Glu132, docking ISG15 onto the Plpro α7 helix (**Extended Data Fig. 5c**). These interactions dislodge the Ubl-fold from the Fingers subdomain (**Figure 2a**). While the complex resembles interaction modes observed in SARS PLpro∼ISG15^CTD^ (pdb 5tl7, ^25^, RMSD of 0.74 Å for PLpro, see **Extended Data Fig. 4**), some interacting residues (especially, Tyr171 on helix α7) are not conserved (**Extended Data Fig. 1, 5c**), and seem to improve the contact in SARS2 PLpro. More variability is seen in MERS PLpro, which binds to ubiquitin and ISG15^CTD^ similarly through its ability to ‘close’ the Fingers subdomain^26^ (see discussion in **Extended Data Fig. 4**).

**Figure 2.**
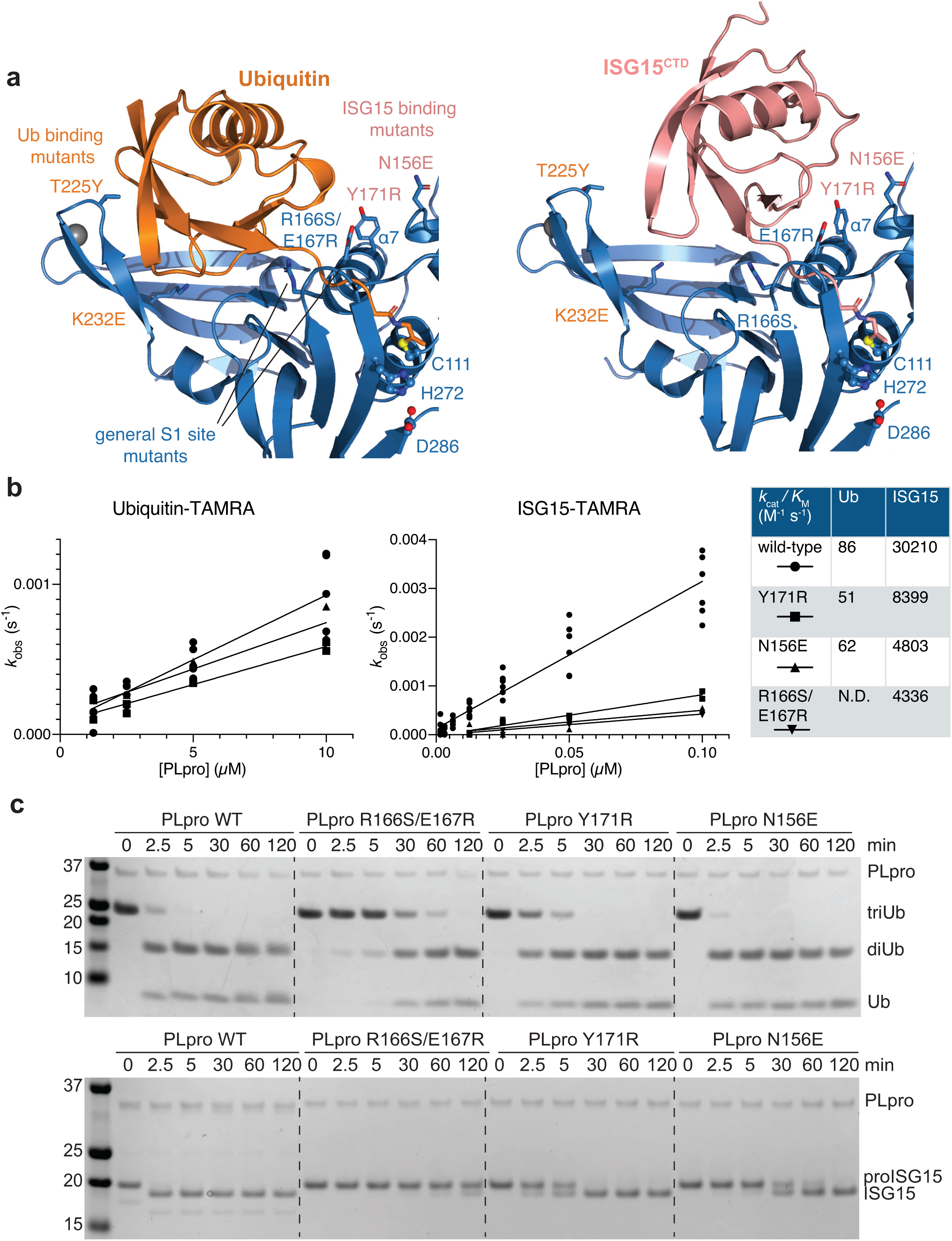
Distinct binding of ubiquitin and ISG15 enables separation of PLpro function. **a**, Detail of the S1 ubiquitin binding site of SARS2 PLpro, bound to ubiquitin (*left*) and ISG15 (*right*), highlighting differential interactions of ubiquitin with the Fingers subdomain, and of ISG15 with the Thumb subdomain of PLpro. Labelled residues were mutated, see **Extended Data Fig. 5. b**, Fluorescence polarisation assays against ubiquitin-TAMRA and ISG15-TAMRA using indicated SARS2 PLpro variants performed in technical triplicate and n=2 for each mutant, and compared to wild-type PLpro as shown in **Extended Data Fig. 2a**. Catalytic efficiencies were calculated as described in **Methods. c**, Gel based analysis of PLpro variant activity against Lys48-triubiquitin and proISG15. Experiments were performed in duplicate, see **Extended Data Fig. 5** and **Supplementary Figure 1**.

Binding mode differences in the S1 ubiquitin binding site provided an opportunity to generate separation-of-function mutations (**Figure 2b,c, Extended Data Fig. 5d-f**). A general S1 site mutant, R166S/E167R ^20^ showed severely diminished activity against either modifier (**Figure 2b,c, Extended Data Fig. 5f**). PLpro N156E (and, to a lesser degree, Y171R) resulted in selective decrease of activity in ISG15 cleavage assays, with little impact on ubiquitin cleavage (**Figure 2b,c, Extended Data Fig. 5f**). Mutations selectively impacting ubiquitin but not ISG15 were more challenging to generate but were apparent by gel-based analysis (see e.g. K232E, **Extended Data Fig. 5f**). Similar experiments have recently been described for MERS PLpro ^27^.

## Impact of the S2 ubiquitin binding site on polyubiquitin and ISG15 cleavage

Polyubiquitin cleavage in SARS2 PLpro is significantly enhanced when a longer ubiquitin chain is used (**Figure 1c, Extended Data Fig. 2b**), due to an S2 ubiquitin binding site provided by the PLpro α2 helix, in which a central Phe residue interacts with the ubiquitin Ile44 patch of the distal ubiquitin in Lys48-diubiquitin (**Extended Data Fig. 1, 6a,b**). Consistently, SARS2 PLpro F69S mutation greatly diminished Lys48-triubiquitin cleavage, without markedly affecting ISG15^CTD^ cleavage (**Extended Data Fig. 6c-f)**. Interestingly and consistent with **Figure 1**, PLpro F69S reduces ISG15-TAMRA hydrolysis ∼3-fold, i.e. to levels observed for PLpro wild-type cleavage of ISG15^CTD^ (compare **Extended Data Fig. 6c, 2a**). A similar trend is observed by gel-based analysis (**Extended Data Fig. 6d-f**). However, in comparison, the effects of S2 site mutations on Lys48-polyubiquitin cleavage (**Extended Data Fig. 6e)**, and of S1 site mutations on ISG15 cleavage (**Figure 2, Extended Data Fig. 5f**), are more pronounced. Mutational analysis hence confirms that the S2 site is more important for providing PLpro with Lys48-polyubiquitin activity and specificity. Taken together, our data illuminate in molecular detail how SARS2 PLpro targets ubiquitin and ISG15.

## Repurposing known drugs to inhibit PLpro activity

We next focussed our attention on the urgent matter of inhibiting PLpro, to confirm its drugability, and to provide new drug candidates with efficacy in treating COVID-19. Ideally, an already clinically approved drug shows a pharmacologically relevant effect on PLpro with sub-µM inhibitory potential, cell penetrance, oral bioavailability, and extensive safety profiles for the required dosage. Such a drug could be expedited for clinical trials.

A 1536-well low-volume high-throughput assay previously used to identify inhibitors of human deubiquitinases (DUBs) ^28^ was adapted for SARS2 PLpro (see **Extended Data Fig. 7a-c** and **Methods**). As a control for complete inhibition, the racemic version of the literature compound 5c ^29^ (here referred to as rac5c, see below), was used at 10 µM concentration and fully inhibited PLpro (**Extended Data Fig. 7d**, orange). A curated library of 5576 compounds, comprising 3727 unique approved drugs and late-stage clinical drug candidates (listed in **Supplementary Information**), was screened in triplicate at 4.2 µM drug concentration (**Figure 3a, Extended Data Fig. 7d-f**).

**Figure 3.**
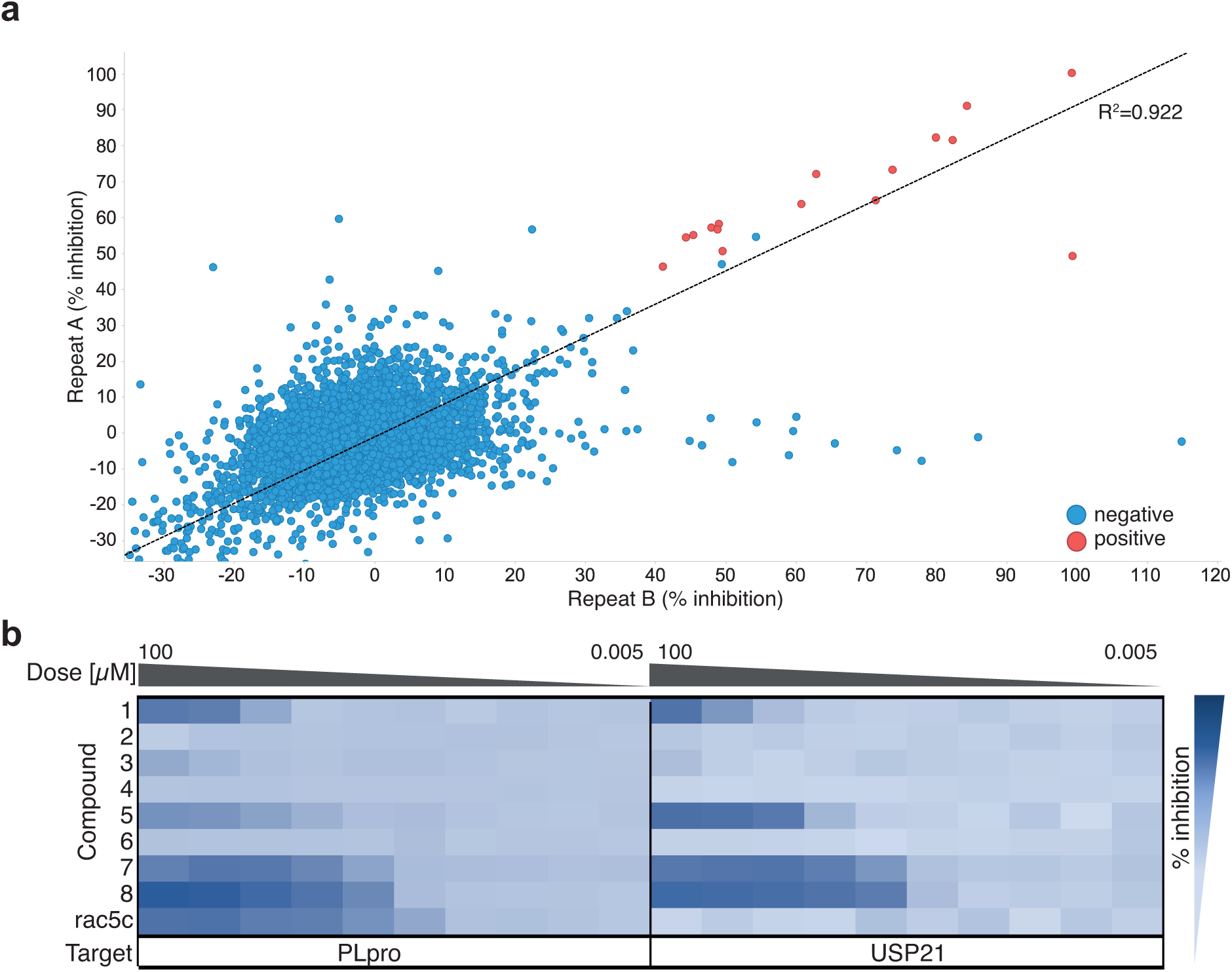
High throughput screen of SARS2 PLpro against known drugs. **a**, High throughput screening of SARS2 PLpro was performed against 5576 approved drugs and late-stage clinical compounds, in 1536-well format using Ub-Rhodamine (see **Methods**). Two replicates out of three are shown; hit compounds were those that inhibited PLpro activity by more than 40% in all three replicates. Correlation (*R*^*2*^) between all screens exceeded 0.89. See **Extended Data Fig. 7** for assay design and quality control, and **Methods. b**, Hit compounds and compound rac5c (see **Figure 4**), were further assessed in 10-point IC_50_ titrations using the Ub-Rhodamine assay, using a starting concentration of 100 µM serially diluted in 1:3 steps. Degree of inhibition is shown as a titration heat-map from dark (full inhibition) to light blue (low/no inhibition). The catalytic domain of human USP21^30^ was used as a counterscreen. Each PLpro hit compound showed either no activity in the titration analysis, or an identical inhibition profile against PLpro and USP21, suggesting assay interference. Rac5c was specific for SARS2 PLpro and did not inhibit USP21 even at the highest concentration of 100 µM. IC_50_ assays were performed in technical triplicate in two independent experiments.

A set of 15 compounds showed 40-90% PLpro inhibition in each triplicate run (**Figure 3a**). Seven of these were excluded as commonly observed false-positives (reactive compounds or dyes that interfere with assays). The remaining 8 compounds were tested in 10-point titration experiments for IC_50_ measurements, as well as counterscreened against the catalytic domain of human USP21 ^30^. We chose this human protein to assess the potential selectivity of inhibition of PLpro over a representative human DUB and as counterscreen. PLpro and USP21 are sufficiently dissimilar to conclude that any compounds inhibiting both with similar IC_50_ would likely be false positives interfering with the assay. After full titration against PLpro and USP21, we found that the 8 hits were either inactive in validation, or equally active towards PLpro and USP21 (**Figure 3b**), suggesting that none of the identified hits are genuine PLpro inhibitors. This contrasted with rac5c, which inhibited PLpro, but did not inhibit USP21 even at 100 µM concentration (**Figure 3b**).

Together, our data suggests that a repurposing strategy using 3727 unique known drugs towards SARS2 PLpro is unlikely to yield drug candidates, and highlights the importance of a counterscreen in assessing the validity of hits coming from a screen of known drugs before any conclusions on their therapeutic potential can be drawn. The robust screen and orthogonal assays for PLpro will be instrumental in drug discovery campaigns.

## Exploiting known SARS PLpro inhibitors against SARS2 PLpro

SARS PLpro has been the focus of academic drug discovery efforts in the last two decades^3^. An initial series of non-covalent small molecules ^31^ was subsequently refined to achieve sub-µM inhibitors of SARS PLpro with high specificity and low cytotoxicity^1,29^. Drug development was aided by structural analysis of several SARS PLpro-compound complexes (**Figure 4a, Extended Data Fig. 8a,b**), showing that compounds bind in the channel occupied by the ubiquitin/ISG15 C-terminal tail, wedged between the SARS PLpro Thumb domain and a so-called Blocking Loop (BL), containing a critical Tyr residue (Tyr269 in SARS, Tyr268 in SARS2) ^1,29^ (**Figure 4a, Extended Data Fig 8a,b**). An extended, Tyr-lacking BL in MERS PLpro (**Extended Data Fig. 1**), renders it unsusceptible to some SARS inhibitors ^32^. Importantly, the BL sequence and length, and all residues involved in inhibitor interactions, are identical between SARS and SARS2 PLpro (**Figure 4a, Extended Data Fig. 1, 8a,b**), suggesting that SARS PLpro inhibitors may have inhibitory potential against SARS2 PLpro.

**Figure 4.**
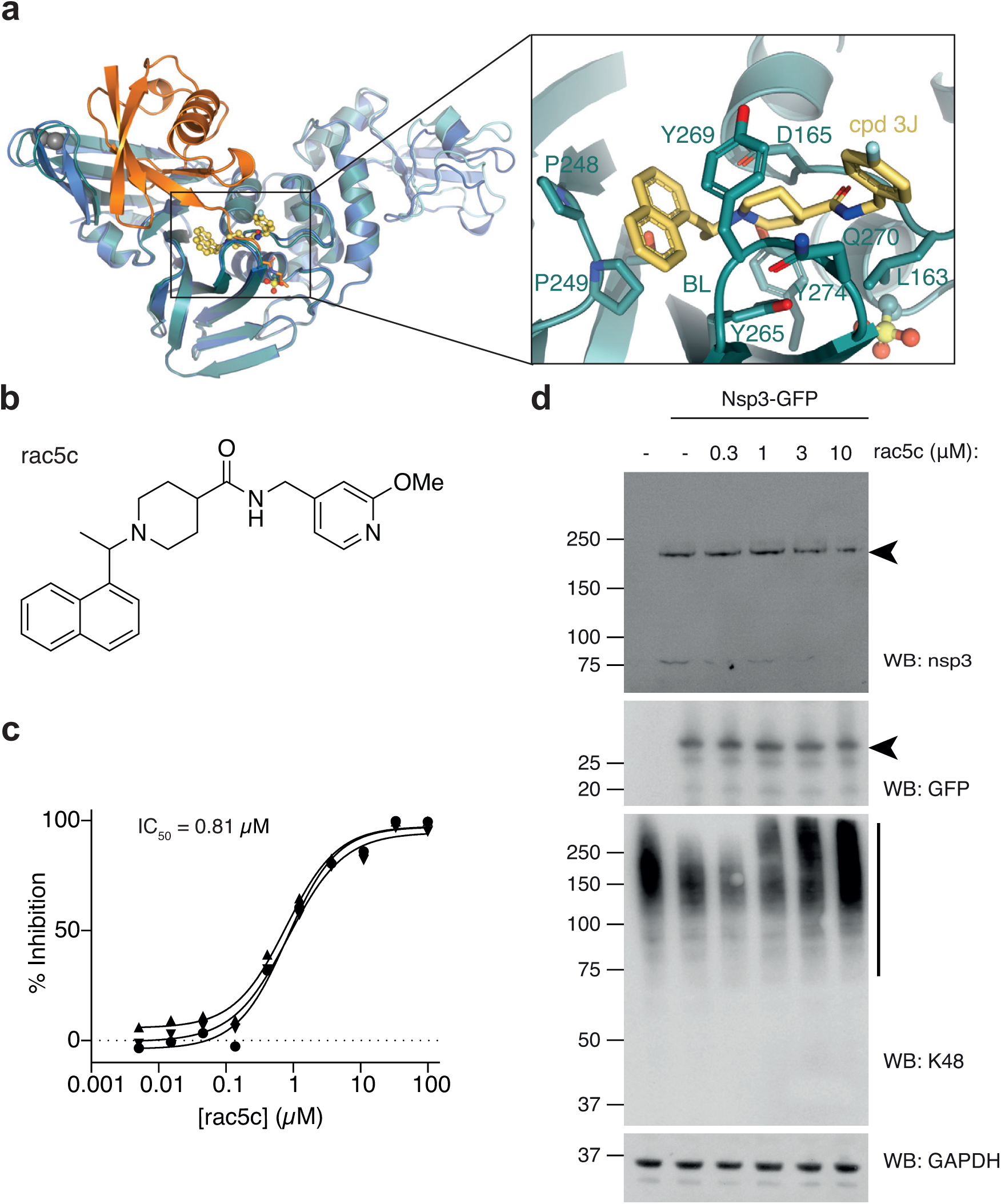
SARS PLpro inhibitors target SARS2 Plpro. **a**, Structure of SARS PLpro bound to compound 3j (cyan/yellow, pdb 4ovz, ^29^) superposed with SARS2 PLpro∼Ub (blue/orange). The inset shows compound 3j bound near the active site. See **Extended Data Fig. 8a,b** for further details. **b**, Chemical structure of rac5c. See **Supplementary Chemistry Methods. c**, *In vitro* inhibition (IC_50_) for rac5c inhibiting SARS2 PLpro. Experiments were performed using the HTS assay (**Figure 3**), in technical triplicate in three independent experiments. A geometric mean was used to determine IC_50_. **d**, Full-length nsp3 was expressed from a C-terminally GFP-tagged vector in HEK293T cells and treated with increasing concentrations of rac5c for 9 h. GFP is released from the C-terminus, presumably by nsp3 activity. Nsp3 can be detected by a SARS2 PLpro antibody (see **Extended Data Fig. 8e** for antibody validation). Lysates were blotted for Lys48-polyubiquitin with a linkage specific antibody. Experiments were performed in duplicate with similar results. Also see **Extended Data Fig. 8f** and **Supplementary Figure 1** for uncropped blots.

We selected and resynthesised racemic forms of three late-stage literature compounds, named according to previous publication ^29^, rac3j, rac3k, and rac5c (see **Supplementary Methods**). IC_50_ measurements performed on our automated screening platform revealed low or sub-µM inhibitory activity for each compound against SARS2 PLpro (**Figure 4b,c, Extended Data Fig. 8c,d**). This confirmed that SARS PLpro inhibitors inhibit SARS2 PLpro.

## SARS2 PLpro inhibitors inhibit nsp3 DUB activity

Nsp3 is a 215 kDa multi-domain enzyme with several catalytic activities. To test whether rac5c would be able to inhibit the PLpro domain in context of full-length nsp3, the protein was transiently expressed from a C-terminally GFP-tagged vector in HEK 293T cells. Full length nsp3 was detected with a SARS2 PLpro specific antibody (validated in **Extended Data Fig 8e**) and its activity was confirmed by proteolytic cleavage of the GFP tag (**Figure 4d, Extended Data Fig 8f**). Nsp3 expression depleted Lys48-linked polyubiquitin, which was inhibited by rac5c in a dose dependent manner (**Figure 4d, Extended Data Fig 8f**). Since it had previously been shown that these inhibitors are specific for PLpro over human DUBs^1^ (also see **Figure 3b**) and since treatment with rac5c did not affect Lys48-linked polyubiquitin in the absence of nsp3 expression (**Extended Data Fig 8f)**, the effect of rac5c on Lys48-polyubiquitin is likely due to inhibition of nsp3/PLpro.

## Antiviral efficacy of SARS2 PLpro inhibitors

All three compounds were tested for their inhibitory potential in Vero monkey kidney epithelial cells infected with SARS-CoV-2. Vero cells undergo extensive cell death upon SARS-CoV-2 infection in contrast to many human cell lines where cytopathic effect (CPE) is less evident^33^. In addition, we found Vero cells were sensitive to DMSO concentrations above 0.3% (v/v), limiting the useful range at which inhibitors could be applied due to their low solubility (**Extended Data Fig. 9a**). Synthesised compounds rac3j, rac3k and rac5e showed no toxicity on Vero cells when used in 0.1% DMSO (enabling compound assessment on cells at concentrations up to 11 µM), but toxicity increased with higher compound and DMSO concentrations (33 µM compound, in 0.3% DMSO) (**Extended Data Fig. 9b**).

Next, compounds were tested in Vero cells infected with SARS-CoV-2 at a multiplicity-of-infection (MOI) of 0.1 (**Figure 5a**), resulting in death of ∼50% of the cell population. Remdesivir (RDV)^34,35^, the only available drug approved for treatment of COVID-19, was used at 12.5 µM concentration^36^, leading to a ∼90% reversal of the SARS-CoV-2 induced CPE. Hydroxychloroquine (HCQ), at 10 µM ^37^, rescued CPE also by ∼90% (**Figure 5b, Extended Data Fig. 9c,d**).

**Figure 5.**
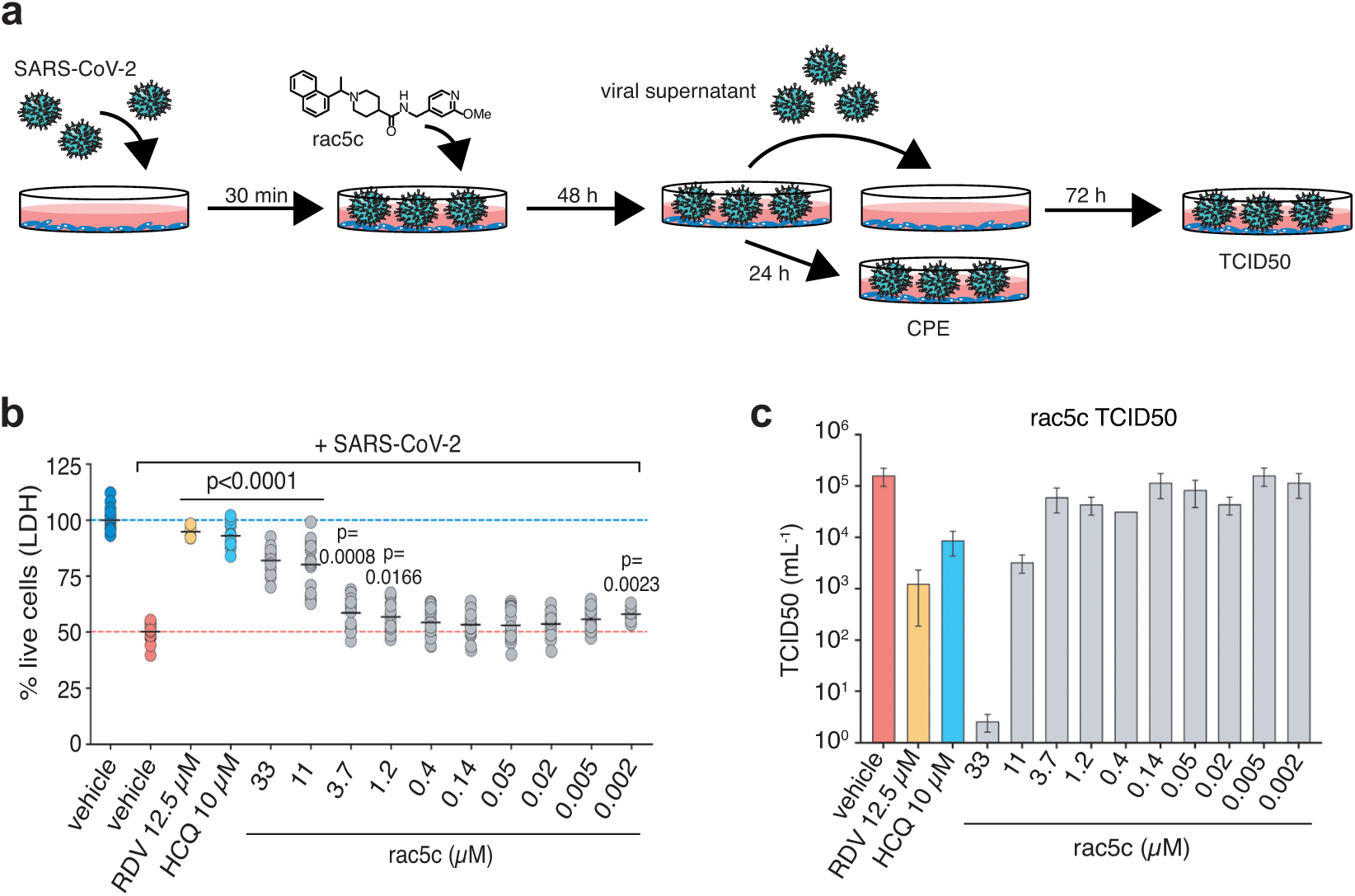
Antiviral effects of SARS2 PLpro inhibitors in an infection model. **a**, Vero cells were infected with SARS-CoV-2 and analysed as shown in the cartoon (see **Methods**). **b**, Reduction in SARS-CoV-2 induced cytopathic effect with rac5c, Remdesivir (RDV) and hydroxychloroquine (HCQ) treatment. DMSO 0.3% (v/v) was required to keep 33 µM rac5c in solution (see **Extended Data Fig 9a,b**). Mean (black line) is provided for 18 samples in each group, representing 3 independent experiments with 6 biological replicates per experiment across the different concentrations of rac5c. HCQ data is pooled from 2 independent experiments and RDV from 1 experiment using 6 biological repeats. P values were calculated using a one-way ANOVA, with regular Dunnet’s post-hoc test for multiple comparisons between treatment arms and infected/vehicle treated control using a single pooled variance. **c**, TCID50 data, mean and SD, for one representative experiment from **b** with 6 technical replicates.

High (33 µM) concentrations of rac5c, rac3j or rac3k, rescued cells from SARS-CoV-2 induced CPE compensating for, and despite, the background toxicity associated with high DMSO concentrations described above **(Figure 5b, Extended Data Fig. 9c,d**). For rac3j and rac3k, this effect diminished at lower concentrations (**Extended Data Fig 9c,d**). For rac5c, treatment at non-cytotoxic concentrations of 11 µM continued to show a marked reduction on CPE, indicating clear antiviral activity (**Figure 5b**).

Antiviral activity is best assessed by a compound’s effect on TCID50 (mean tissue culture infection dose) in which cell supernatant from infected cells is assessed for infectious viral titre in secondary infections. RDV (12.5 µM) and HCQ (10 µM) reduced viral titre by 100- and 10-fold, respectively. SARS2 PLpro inhibitors, at concentrations that protected cells from CPE, showed a 3-4 log decrease in infectious viral titre at 33 µM (**Figure 5c, Extended Data Fig. 9e,f**), and rac5c at 11 µM decreased viral titre to a similar extended as RDV and HCQ treatment. In a second model system, the human lung adenocarcinoma cell line Calu-3, infection with SARS-CoV-2 decreased Lys48-polyubiquitin levels, which could be rescued with rac5c treatment (**Extended Data Fig. 9g, h**). Together, our data highlight that inhibition of SARS2 PLpro with small molecules can have striking antiviral effects.

## Conclusion

The biochemical activities and structural properties of the PLpro domain of the essential SARS-CoV-2 protein nsp3 hold tremendous promise as a target to generate a new class of antivirals for coronaviruses. The three distinct substrates of PLpro, namely the viral polyprotein, degradative Lys48-polyubiquitin and antiviral ISG15 signals, make PLpro an excellent candidate for pharmacological intervention. Detailed molecular understanding of how PLpro targets ubiquitin and ISG15, and a robust high-throughput screen, pave the way to structure-guided drug discovery. Indeed, while available clinically tested drugs may not be suitable to target PLpro (**Figure 3**), exciting pharmacologically unrefined lead compounds are already available to specifically target SARS2 PLpro. We show that these compounds have antiviral efficacy and seem as potent as other drugs that target viral replication (e.g. the viral polymerase inhibitor Remdesivir). Moreover, the direct antiviral effect of PLpro inhibitors is likely further supplemented by suppressing PLpro’s role in subverting the innate immune system. PLpro inhibitors may prove useful in restarting (and/or rebalancing) host processes pathologically deregulated in COVID-19.

## Supporting information

Screening Table

Uncropped Gels

Chemistry Methods

Screening Results

## Acknowledgments

We would like to thank Kanta Subbarao (Peter Doherty Institute, Melbourne) for live SARS-CoV-2 virus, Frederick P. (Fritz) Roth (University of Toronto) for providing the SARS2 nsp3 pENTRY plasmid, Thibault Major (University of British Columbia, Vancouver) for vectors and for supporting J. Bernardini, Sandra Nicholson, Peter Colman, Ian Wicks and Thomas Cotton (WEHI) for sharing expertise and reagents, and Jonathan O’Connell (FORMA Therapeutics) for advice on high-throughput assays.

This work was funded by The Walter and Eliza Hall Institute of Medical Research, an NHMRC/MRFF ‘VirDUB’ grant MRF2002119 (to DK, GL, MP, PEC), NHMRC Investigator Grants and Fellowships (GNT1178122 to DK, GNT0637350 to MP, and GNT1117089 to GL), NHMRC Independent Research Institutes Infrastructure Support Scheme grant (361646) and Victorian State Government Operational Infrastructure Support grant, and a generous donation by Hengyi Pacific Pty Ltd to support COVID-19 research.

Screening was conducted at the Walter and Eliza Hall Institute’s National Drug Discovery Centre (NDDC). The NDDC received grant funding from the Australian Government and the Victorian State Government, with additional support from generous philanthropic donors including Mike Fitzpatrick, Helen Sykes and AWM Electrical. WEHI’s screening facilities are also supported by Therapeutic Innovation Australia (TIA). TIA is supported by the Australian Government through the National Collaborative Research Infrastructure Strategy (NCRIS) program. The Australian Drug Discovery Library (ADDL) was compiled with the financial assistance of MTPConnect. We acknowledge Compounds Australia (www.compoundsaustralia.com) for their provision of specialised compound management and logistics services to the project. This research was undertaken in part using the MX2 beamline at the Australian Synchrotron, part of ANSTO, and made use of the Australian Cancer Research Foundation (ACRF) detector.

## Author contributions

DK, GL, MP and PEC conceived the project and obtained funding. TK produced pure PLpro, performed FP assays, crystallised PLpro bound to ubiquitin, and determined structures. Protein production, gel-based activity assays, and probe generation was performed by DJC and LWR. YS and ZYG generated and purified mutant PLpro. PPG and HO contributed ISG15^CTD^-TAMRA reagent. DJC crystallised the PLpro∼ISG15^CTD^ complex with help from LWR. JN set up protein crystallisation and guided crystal optimisation. TK, BCL and DK collected synchrotron data with support from AR-T and PEC, and TK and BCL refined structures. NWK, JPM, CG, and GL designed and performed compound synthesis. BGCL and KNL designed and established high throughput assays, and BGCL, AEA, TRB, JPM and KNL performed high throughput screens and analysed data. JPB, GD, RF designed and performed nsp3 and PLpro expression studies in cells. GE, CCA, JPC, MD and MP designed and established SARS-CoV-2 infection models and analysed data. DK wrote the manuscript with help from all authors.

## Author information

The authors declare no competing interests. All reagents and materials are available upon reasonable request from the corresponding author.

## Online Methods

### Molecular Biology

#### Generation of bacterial expression vectors

The sequence of SARS CoV-2 PLpro (amino acids (aa) 1563-1878, with aa E1564 designated as residue 1, according to previously published numbering) was based on the polyprotein orf1ab (GeneBank: QHD43415) and was purchased as a codon-optimised gene-block (IDT) for bacterial expression. PLpro was cloned by ligation independent cloning into the pOPIN-B vector ^38^ using the In-Fusion HD cloning Kit (Takara Clontech). All PLpro mutants were introduced by site directed mutagenesis using the Q5 Site-Directed Mutagenesis Kit (NEB).

For Ub-PA and ISG15^CTD^-PA preparation, Ub (1-75) and ISG15^CTD^ (79-156) genes were expressed from pTXB1 vectors as described^22,39^.

proISG15 (2-165) and proISG15^CTD^ (79-165) were expressed from pOPIN-B vectors as described in ^22^.

#### Generation of mammalian expression vectors

For mammalian expression, SARS2 PLpro variants, SARS PLpro (aa 1541-1855 of polyprotein 1ab) and MERS PLpro (aa 1482-1803 of polyprotein 1ab) were generated as codon optimised gene-block (IDT) for bacterial expression, and were transferred into pOPIN-F using In-Fusion HD cloning (Takara Clontech).

Full length nsp3 (kindly provided by Fritz Roth, University of Toronto) was cloned from a pENTRY vector into the pDEST47 vector using gateway cloning with the LR clonase mix (Invitrogen), according to manufacturer’s instructions.

### Protein purification

#### PLpro

PLpro wild-type and mutant expression vectors were transformed into *E. coli* Rosetta 2(DE3) pLacI competent cells (Novagen) and bacterial cells were grown in 2xYT medium at 37 °C. At OD_600_ = 0.8 the temperature was reduced to 18 °C and expression was induced with 0.3 mM IPTG. Cells were harvested 16 h post induction and stored at -20 °C.

For purification, cells were resuspended in lysis buffer (50 mM Tris pH 7.5, 300 mM NaCl) supplemented with lysozyme, DNaseI and cOmplete EDTA-free protease inhibitor cocktail tablets (Roche) and lysed by sonication. Lysates were cleared by centrifugation at 40,000 *g* for 30 min at 4 °C and His-tagged proteins were purified either by using a HisTrap FF column (5 mL, Cytvia) with gradient elution over 5 column volume (CV) from buffer A (20 mM Tris pH 7.5, 300 mM NaCl and 10 mM Imidazole) to buffer B (20 mM Tris pH 7.5, 300 mM NaCl and 400 mM Imidazole), or with Ni-NTA HisBind resin (EMD Millipore) eluting with buffer B (2x 10 mL). Pooled fractions were supplemented with His-3C protease for His-tag cleavage and dialysed overnight at 4 °C (for wt: 50 mM Tris pH 7.5, 300 mM NaCl, 5 mM β-mercaptoethanol (bME), for mutants: 20 mM HEPES pH 7.5, 300 mM NaCl and 10 mM bME). His-3C protease and His tags were removed by Ni-NTA HisBind resin (EMD Millipore) and proteins were further purified by size exclusion chromatography using a HiLoad 16/600 Superdex 75 pg column (GE Healthcare) equilibrated with storage buffer (20 mM HEPES pH 7.5, 150 mM NaCl, 1 mM TCEP). Protein samples were concentrated, flash frozen in liquid nitrogen and stored at -80 °C.

#### Thermal shift assay

Thermal shift assays were performed for quality control after PLpro wt and mutant purification, using the Tycho NT.6 (NanoTemper Technologies). PLpro wt and mutants were measured at 1 μM in storage buffer. The inflection temperatures of each protein were calculated by the Tycho NT.6 software (1.2.0.750). Technical duplicates were measured in two independent experiments. Data were analysed using GrapPad Prism.

#### His_6_-proISG15 and His_6_-proISG15^CTD^ purification

proISG15 and proISG15^CTD^ were expressed as described above but induced with 0.2 mM IPTG and resuspended in buffer C (50 mM Tris pH 7.5, 150 mM NaCl, 2 mM bME) prior storage at -20°C. Affinity purification was performed as for PLpro but with the following modifications. HisTrap FF resin was washed with buffer C supplemented with 15 mM imidazole and eluted with a linear gradient of 10 CV from buffer C to buffer D (buffer C supplemented with 300 mM imidazole). Eluted proteins were diluted 10-fold to a low salt buffer (50 mM Tris pH 8.0, 30 mM NaCl, 2 mM bME) and passed over a ResourceQ column (Cytvia). The eluted proteins were concentrated and further purified by size exclusion chromatography (HiLoad 16/600 Superdex 75pg, Cytvia) into Buffer C. Protein containing fractions were concentrated, flash frozen and stored at -80°C until further use.

#### Generation of ubiquitin and ISG15^CTD^ suicide probes

Ub-intein and ISG15^CTD^-intein proteins were expressed as for PLpro. Cell pellets were resuspended in Buffer E (20 mM HEPES, 50 mM NaOAc pH 6.5, 75 mM NaCl) and Buffer F (50 mM HEPES, 100 mM NaOAc pH 6.5) respectively.

Ub-MesNa (2-mercaptoethansulfonate as a sodium salt) and subsequently Ub-PA were prepared as described previously^39-41^. Human ISG15^CTD^ (79-156)-MesNa (ISG15^CTD^-MesNa) and the ISG15^CTD^-PA suicide probe were prepared as described previously^42^.

The completed reactions underwent final size exclusion chromatography (HiLoad 16/600 Superdex 75pg, Cytvia) into Buffer E (Ub-PA) or Buffer F (matISG15^CTD^-PA). The resultant fractions were concentrated, flash frozen and stored at -80°C until further use.

#### Preparation of the PLpro∼Ub and PLpro-ISG15^CTD^ complex for crystallisation

Purified PLpro was incubated with 3x molar excess of either Ub-PA or ISG15^CTD^-PA at RT for 2 h. Unreacted probe was separated from the complex by size exclusion chromatography (Superdex 75 Increase 10/300 GL) into 20 mM Tris pH 7.5, 150 mM NaCl, 1mM TCEP (PLpro∼Ub) or 20 mM HEPES pH 7.5, 150 mM NaCl, 1 mM TCEP (PLpro∼ISG15^CTD^) and the eluted complexes were concentrated to 4 mg/mL for PLpro∼Ub and 5 mg/mL or 8 mg/mL for PLpro∼ISG^CTD^ for crystallisation.

#### Crystallisation

Crystallisation screening was performed at the CSIRO’s Collaborative Crystallisation Centre (C3) in Melbourne, Australia. For PLpro∼Ub at 4 mg/mL, one crystal grew in 30% (w/v) PEG 4000, 0.2 M sodium acetate, 0.1 M Tris chloride pH 8.5, in a 96-well, sitting drop vapour diffusion plate (150 nL protein to 150 nL reservoir solution) at 20 °C. The crystal was cryoprotected with 20% (w/v) PEG 4000, 0.2 M sodium acetate, 0.1 M Tris chloride pH 8.5 and 25% (v/v) glycerol before vitrification in liquid nitrogen. For PLpro∼ISG15^CTD^ initial crystals grew at concentrations of both 5 mg/mL and 8 mg/mL complex, in several conditions containing 0.2 M lithium or ammonium sulfate, 25% (w/v) PEG 3350 and bis-tris chloride pH 5.5 to 6.5, at 20 °C. The structure was solved from a crystal reproduced in a hanging drop 24-well plate using 5 mg/mL protein complex grown in 0.2 M lithium sulfate, 25% (w/v) PEG 3350 and 0.1 M bis-tris chloride pH 6.5 and a protein to reservoir ratio of 1 µL to 0.5 µL. The crystal was stepwise cryoprotected by using the mother liquor supplemented with 15% (v/v) glycerol as a first and 28% (v/v) glycerol as a second step, before vitrification in liquid nitrogen.

#### Data collection, phasing and refinement

Diffraction data were collected at the Australian Synchrotron (Australian Nuclear Science and Technology Organisation, ANSTO) beamline MX2^43^ (wavelength: 0.953725 Å, temperature: 100K). Collected datasets were processed and scaled with XDS ^44^ and Aimless ^45^ (within CCP4suite^46^). The structures of SARS2 PLpro∼Ub and SARS2 PLpro∼ISG15^CTD^ were solved by molecular replacement to a resolution of 2.7 and 2.9 Å respectively, using Phaser ^47^ and the apo structure of SARS2 PLpro (pdb: 6wrh, unpublished) and either ubiquitin (from pdb 5ohk, ^39^) or ISG15^CTD^ (from pdb 6ffa, ^22^) as search models.

Refinement and model building was performed in PHENIX ^48^ and Coot^49^. Both structures were initially refined by cartesian stimulated annealing following rigid body refinement. For both complexes, secondary structure restrains were set and the apo structure of SARS2 PLpro (pdb 6wrh, unpublished) was used as reference model. TLS parameters were set to one TLS group per chain. For SARS2 PLpro∼ISG^CTD^ additional NCS refinement was utilised in each refinement cycle. For the covalent linkage of the propargylamide to the catalytic Cys111 of SARS2 PLpro in each structure, geometric restrains for propargylamide (AYE) derived from PHENIX elbow and a parameter file defining the linkages was used. Models were validated using MolProbity ^50^ and Coot indicating for PLpro∼Ub following Ramachandran plot statistics: 0.0% outliers, 2.63% allowed and 97.37% favoured and for PLpro∼ISG^CTD^: 0.0% outliers, 2.60% allowed, 97.40% favoured.

Structural figures were generated using PyMol. Further data collection and refinement statistics can be found in **Extended Data Table 1**.

### PLpro activity assays

#### Gel-based PLpro DUB activity and chain specificity assays

Gel-based cleavage assays were performed as previously described ^51^ with the following modifications. Reactions were initiated at room temperature (23°C) in a final volume of 150 μL (for the specificity assay) or 350 μL (for longer time-course assays) and 20 mM Tris pH 7.5, 100 mM NaCl, 10 mM DTT was used as the reaction buffer. Triubiquitin substrates were enzymatically assembled as previously described ^52^. Final enzyme and substrate concentrations were 0.25 μM and 2 μM respectively. Reactions were stopped at indicated time points by mixing 20 μL of reaction with 20 μL 2x NuPAGE LDS sample buffer (Invitrogen) and analysed by SDS-PAGE (Invitrogen NuPAGE™ 4-12 % Bis-Tris) and Coomassie staining (Instant Blue, Expedeon).

For gel-based quantitative analysis, Coomassie stained gel images were converted to grayscale and band intensities were quantified using ImageLab™ (Bio-Rad). Background intensities were automatically subtracted using a base line relative to the lowest contrasting band for each gel. Values were then normalised to the PLpro band in each lane. Remaining substrate concentrations were calculated with respect to the substrate concentration at time point zero (100%). The resulting values were plotted over time and the initial values within the linear range were used to calculate the relative activity measures.

#### Fluorescence polarisation (FP)-based PLpro activity assays

FP-assays were performed with Ub-KG-TAMRA (UbiQ bio), mouse ISG15-KG-TAMRA (UbiQ bio) and ISG15^CTD^ -TAMRA ^22^ to determine the catalytic efficiencies for PLpro wt and mutants. For the assay small volume, non-binding, black bottom, 384-well plates were used, and reactions were measured on a CLARIOstar plus plate reader (BMG Labtech) using optical settings for the TAMRA fluorophore (excitation: 540 nm, emission: 590 nm). Before each measurement the instrument settings were referenced to 50 mP KG-TAMRA control at a concentration of 50 nM. All substrates were used at a final concentration of 150 nM, while the dilution series of the enzyme concentrations varied according to the substrate (10 µM - 0.156 µM for Ub-cleavage; 100 nM - 1.56 nM for ISG15- and ISG15^CTD^-cleavage; 250-3.90 nM for ISG15^CTD^-cleavage to determine the catalytic efficiency). Enzyme (SARS2 PLpro wild-type and mutants) and substrates were diluted in assay buffer (20mM HEPES pH 7.5, 150mM NaCl, 1mM TCEP, 50 µg/mL BSA) to 2x concentrations and reactions were started upon addition of 2x enzyme to 2x substrate in a final volume of 15 µL. Kinetics were measured in technical triplicates over 60 minutes with one read per minute in at least two independent experiments with the exact number indicated in the figure legends. For the determination of the catalytic efficiency of SARS2 PLpro wild-type on ISG15^CTD^, two of the four independent measurements were performed in technical duplicates, due to substrate limitations.

Data were analysed using the CLARIOstar software MARS, Microsoft Excel and GraphPad Prism (version 8.3.1). Measured fluorescence polarisation values were blank corrected (with a buffer only control) and converted into anisotropy (mA) using the CLARIOstar MARS software. Technical replicates were averaged and fitted by non-linear curve fitting using one-phase decay in GraphPad Prism. The determined rate constants (k_obs_) were then plotted over the enzyme concentrations and fitted using linear regression, to determine the catalytic efficiency *k*_cat_/*K*_M_ as the slope.

### High throughput screening (HTS)

#### Ub-Rhodamine PLpro activity assays for HTS

For HTS screening, PLpro activity was monitored in a homogenous fluorescence intensity assay using the substrate Ub-Rhodamine110Gly (UbiQ bio, here referred to as Ub-Rhodamine). Experiments were performed in either 384-well or 1536-well black non-binding plates (Greiner 784900 and 782900 respectively) with a final reaction volume of 6 µL. The assay buffer contained 20 mM Tris (pH 8), 1 mM TCEP, 0.03% (w/v) BSA and 0.01% (v/v) Triton-X.

PLpro at a final concentration of 50 nM, was added to the plates (preparation of screening plates described below) and incubated at room temperature for 10 min. Ub-Rhodamine (final concentration 100 nM) was added to start the reaction and incubated for 12 min at room temperature. For end-point assays the reaction was stopped by the addition of aqueous citric acid (1 µL) at a final concentration of 10 mM. All reagents were dispensed using the CERTUS FLEX (v2.0.1, Gyger) and Microplates were centrifuged using a Microplate Centrifuge (Agilent). The reaction was monitored by an increase in fluorescence (excitation 485 nm and emission 520 nm) on a PHERAstar® (v5.41, BMG Labtech) using the FI 485 520 optic module. The HTS screen was performed with one measurement for each compound in three independent experiments.

In the counterscreen the deubiquitinating enzyme USP21 (final concentration 5 nM) was used within the same setting, but using an incubation time of 2 min after addition of UbRhodamine110, before the reaction was stopped. Counter and confirmation screen were performed with 3-6 technical replicates in two independent experiments.

#### Screen preparation and data analysis

We assessed the activity of 5,577 compounds contained in commercially available libraries of known drugs (Sigma-Aldrich LOPAC, Tocris and Prestwick) as well as in-house curated collections of FDA approved drugs and advanced pre-clinical compounds (for a complete compound list, see **Supplementary Information**). Analysis of these libraries identified 3727 unique compounds. Compounds were obtained from Compounds Australia, where they are stored under robust environmental conditions.

Assay-ready plates were prepared by dry-spotting compounds in DMSO using an Echo® Acoustic Dispenser (LabCyte). Compounds were tested at 4.2 µM in final 2% (v/v) DMSO. The screen was run using instruments integrated with Momentum Laboratory Automation software (v5.3.1, Thermo Fisher Scientific). Data was normalised to 2% (v/v) DMSO (negative control, 0% inhibition) and 100 µM rac5c (positive control, 100% inhibition). Screen assay quality was monitored by calculation of robust Z’ by the following formula where (+) denotes the positive controls (low signal), (-) denotes the negative controls (high signal) and MAD is the median absolute deviation:

robust Z’ = 1-(3*(MAD-+ MAD^+^) / abs(median--median^+^))

where MAD = 1.4826 * median(abs(x – median(x)))

Plates were excluded from analysis if robust Z’ < 0.5. Hits were selected as > 4*MAD over the median of the negative control.

To determine the potency of the inhibitors, a series of 10-point, 1:3 serial dilutions was performed from a highest starting concentration of 100 µM. The 10-point titration curves were fitted with a 4-parameter logistic nonlinear regression model and the IC_50_ reported is the inflection point of the curve. Data were analysed in TIBCO Spotfire® 7.11.2.

### Chemical synthesis of rac3k, rac3j and rac5c

The chemical synthesis, purification and characterisation of compounds are described in the **Supplementary Information**.

### Cell-based studies and infection assays

#### Cell lines used

HEK293T, Vero (CCL-81) and Calu-3 cells displayed expected cell morphologies and were sent for validation to Garvan Molecular Genetics facility (on 15 June 2020). Cell lines were screened on a monthly basis for mycoplasma contamination using the PlasmoTest kit (Invivogen) as per manufacturer’s instructions. All used cells were mycoplasma free.

#### Cell culture

For infection studies, Vero (CCL-81) cells were cultured in Dulbecco’s Modified Eagle Medium (DMEM + 1g/L D-Glucose, L-Glutamine and 110 mg/L Sodium Pyruvate; Gibco) supplemented with 10% (v/v) heat-inactivated fetal bovine serum (FBS; Sigma-Aldrich), 100 U/mL penicillin and 100 mg/mL streptomycin at 37°C and 5% CO_2_. Vero cells were seeded in a volume of 100 µL DMEM medium into tissue culture treated flat-bottom 96-well plates (Falcon) at a density of 1 × 10_4_ cells/well and incubated over night before infection and/or treatment at confluency.

HEK293T cells were cultured DMEM with 10% (v/v) FBS (Gibco), penicillin (100 U/mL) and streptomycin (100 µg/mL) at 37°C with 10% CO_2_. Cells were seeded in 6 well or 24 well plates and transfected with pOPINF vectors encoding MERS PLpro, SARS PLpro, SARS2 CoV-2 PLpro or a pDEST47 vector encoding nsp3-GFP when cells were at 70-80% confluency with Lipofectamine 3000 (Invitrogen) as per manufacturer’s instructions. 48 h post transfection, cells were harvested for immunoblotting.

#### Cytotoxicity and antiviral efficacy by LDH release cell death assay

Viability of uninfected and vehicle (DMSO) or Bcl2-inhibitor ABT-199 and Mcl-1 inhibitor S63845 treated, or uninfected and SARS-CoV-2-infected and/or Plpro inhibitor treated Vero cells was determined using the CytoTox 96^®^ Non-Radioactive Cytotoxicity Assay (Promega) 72 hours post infection/treatment. The percentage of living cells was calculated comparing LDH release of surviving cells in infected and/or treated cells to LDH release of non-infected or non-treated control cells. Prism 8 software (GraphPad) was used to perform statistical tests in **Figure 5** and **Extended Date Fig.9**. Groups were compared as stated in figure legends.

#### SARS-CoV-2 infection and inhibitor treatment

SARS-CoV-2 was obtained from The Peter Doherty Institute for Infection and Immunity (Melbourne, Australia), where the virus was isolated from a traveller from Wuhan arriving in Melbourne and admitted to hospital in early 2020. Viral material was used to inoculate Vero/hSLAM cells for culture, characterisation and rapid sharing of the isolate^53^.

Vero cells were seeded and rested overnight to confluency in flat-bottom 96-well plates and washed twice with serum free DMEM medium and infected with SARS-COV-2 and MOI of 0.1 or 1 in 25 ul of serum free medium. Cells were cultured at 37°C and 5% CO_2_ for 30 minutes. Cells were topped up with 150 µL of serum free medium containing PLpro inhibitor compounds at various concentrations in 6 replicates per concentration. Cells were monitored daily by light microscopy for morphological changes resulting from virus cytopathic effect. Viability of cells was assessed at day 3 post infection / treatment by LDH release cell death assay as described above.

#### Median Tissue Culture Infectious Dose (TCID50) assay

Vero cell culture supernatant of SARS-CoV-2 infection / treatment assays were harvested 2 days after infection / treatment and diluted in 5x 1:7 serial dilutions in a round-bottom 96 well plate (Falcon) and 6 replicates per dilution. Vero cells were seeded and rested overnight to confluency in flat-bottom 96-well plates and washed twice with serum free DMEM medium. 25 µL of serially diluted virus was added onto washed cells and cultured at 37°C and 5% CO_2_ for 30 minutes before cells were topped up with 150 µL of serum free medium. Cells were monitored at day 2 post infection / treatment by light microscopy for morphological changes resulting from virus cytopathic effect. Virus concentration where 50% of cells show CPE in comparison to untreated cells was defined as TCID50 factor.

The TCID50 calculation is performed using the Spearman and Kärber method, which provides the mean and standard deviation after scoring 300 wells per drug (CPE or not) across the range of dilutions.

#### SARS-CoV-2 RNA extraction from Calu-3 cells and quantitative real time PCR

SARS-CoV-2-infected Calu-3 cells were lysed and RNA extracted using the RNeasy 96 kit (Qiagen). Total cell lysate RNA was reverse transcribed and amplified on the LightCycler 96 (Roche) using the Taqman RNA-to-Ct 1-step kit (Applied Biosystems) to detect virus using the specific SARS-CoV-2 N1 primer/probe assay (IDT). Data was analysed on LightCycler 96 software (Roche).

#### Immunoblotting

Lysates were generated by lysis in 50 mM Tris-Cl, 150 mM NaCl, 1% (v/v) NP-40 with complete protease inhibitors (Roche) and quantified by BCA assay as described. SDS-PAGE was performed with between 20 and 70 ug of protein lysate run per well. Following SDS-PAGE, gels were transferred to 0.2 µm PVDF membranes using the Transblot Turbo system (BioRad). Membranes were blocked in 5% (w/v) milk powder in Tris buffered saline with 0.05% (v/v) Tween-20 (Sigma, TBS-T) for 1 h and then incubated overnight in primary antibody diluted in 5% (v/v) BSA in TBS-T (PLpro antibody, chicken polyclonal, 1:250, (Lifesensors, #AB-0602-0250); anti-Ubiquitin antibody Lys48-specific (Apu2), rabbit monoclonal, 1:1000, (Sigma Aldrich, #05-1307); GAPDH mouse monoclonal antibody (6C5), 1:3000, (Invitrogen, #AM4300); anti-GFP antibody chicken polyclonal, 1:1000, (Abcam, #ab13970)). Following 3 TBS-T washes, membranes were incubated with conjugated secondary antibodies in TBS-T for 1 h at room temperature (IRDye 800CW goat anti-mouse IgG secondary, 1:10000 (Li-Cor, #925-32210); goat anti-chicken IgY-HRP, 1:10000 (SantaCruz, #sc-2428); rabbit IgG HRP, 1:10000, (GE Healthcare, # NA934VS)). Following an additional 3 washes in TBS-T, membranes were developed with fluorescence detection or with Clarity Western ECL chemiluminescence substrate (BioRad) as per manufacturer’s instructions using the Chemidoc (BioRad).

### Data availability statement

Coordinates and structure factors have been deposited with the protein data bank under accession codes 6xaa, 6xa9. Uncropped versions of all gels are displayed in **Supplementary Figure 1**. All reagents and data are available upon reasonable request from the corresponding author.

**Extended Data Figure 1.**
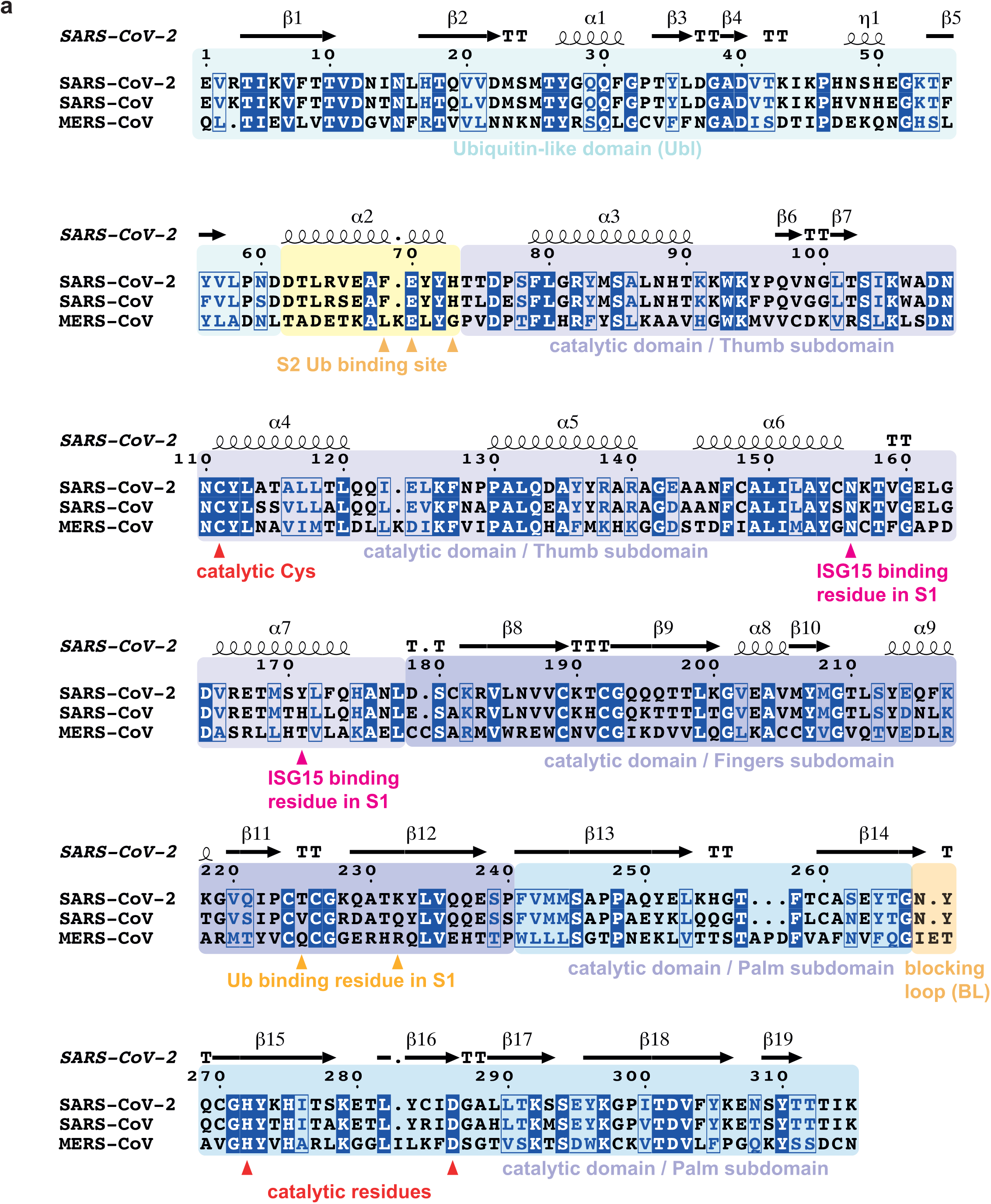
Annotated sequence alignment for Coronavirus PLpro. **a**, Sequence alignment generated with T-coffee/ESPRIPT ^54,55^ aligning PLpro sequences from SARS2, SARS and MERS. Sequence numbering and secondary structure elements are shown according to the high-resolution apo structure of SARS-CoV-2 PLpro (pdb 6wrh, unpublished). T, turn. Domains and subdomains are boxed in different colours, and catalytic triad, as well as residues mutated in this study, are indicated.

**Extended Data Figure 2.**
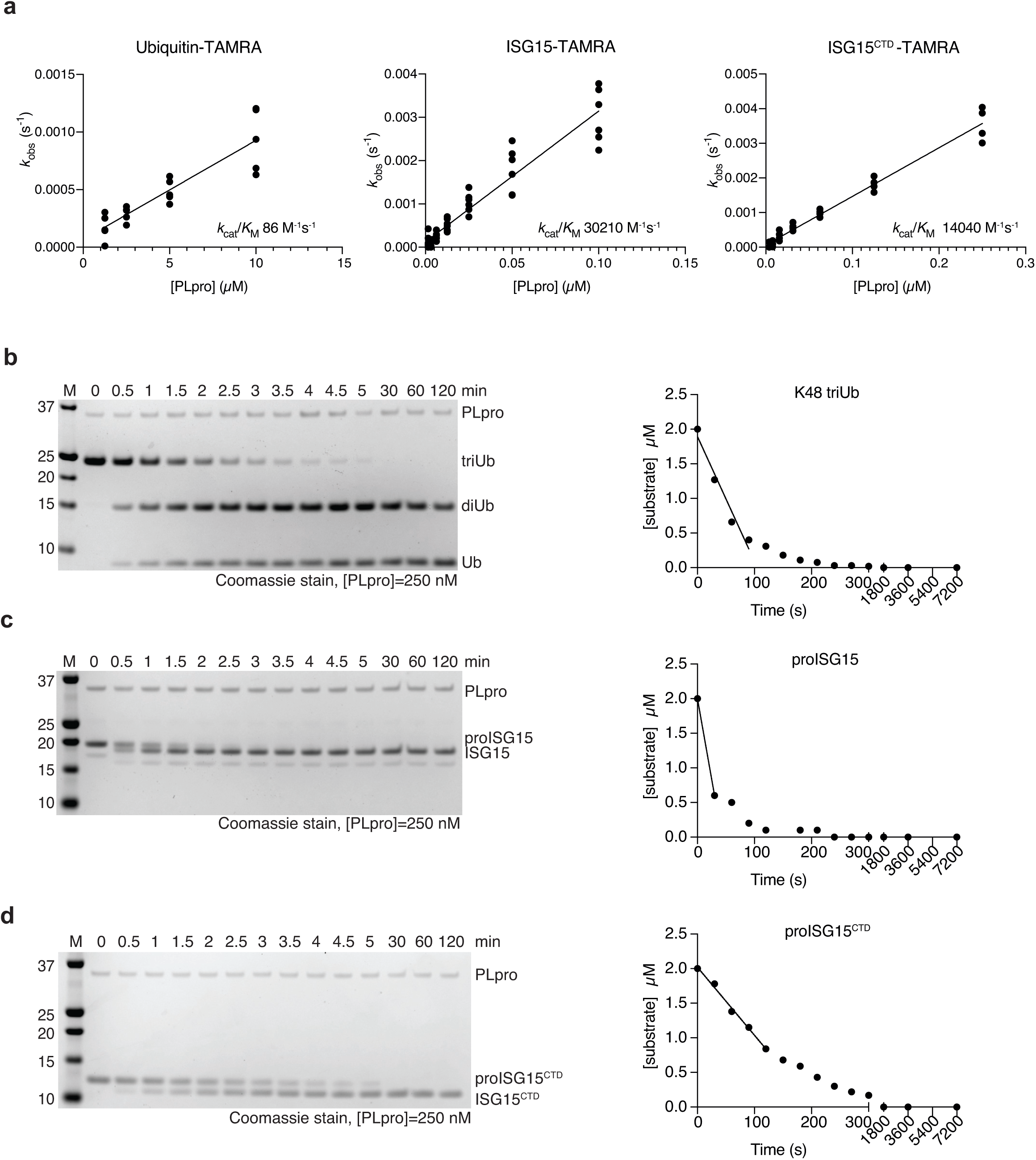
Gel-based kinetics and catalytic efficiency. **a**, Raw data of **Figure 1b** was used to calculate catalytic efficiency (*k*_cat_*/K*_M_). Experiments were performed in technical triplicate and n=5 (Ub-TAMRA), n= 6 (ISG15-TAMRA), n=4 (ISG15^CTD^-TAMRA) biological replicates. See **Methods** for further details. **b**, A monoubiquitin-based substrate, such as Ub-TAMRA, is an inefficient substrate for PLpro as the enzyme prefers longer Lys48-linked ubiquitin chains, which it binds via S1 and S2 sites^20^. Hydrolysis of triubiquitin into mono- and diubiquitin, hence enables a better estimation of the true ability of PLpro to target polyubiquitin (see **Figure 1c**). **b**, In order to quantify SARS2 PLpro activity, triubiquitin cleavage was followed over a time course, resolved on Coomassie-stained SDS-PAGE gels. The disappearance of triubiquitin was quantified by densitometry, plotted to the right. The linear part of the data, corresponding to an estimated observed catalytic rate (*k*_obs_), is indicated by a line. **c, d**, For direct comparison, hydrolysis of extended ‘pro’-forms of human ISG15, in which the modifier is extended by 8 residues on the C-terminus, to the mature form sporting a free Gly156-Gly157 C-terminus, was analysed as in **b**, as previously described^22^. The loss of 8 residues can be visualised as a small shift in size by SDS-PAGE. Disappearance of the higher molecular weight band was quantified by densitometry and plotted on the right. The linear part of the curve was used to visually indicate *k*_obs_. Experiments were performed in duplicate and quantified as shown in **Supplementary Figure 1**.

**Extended Data Figure 3.**
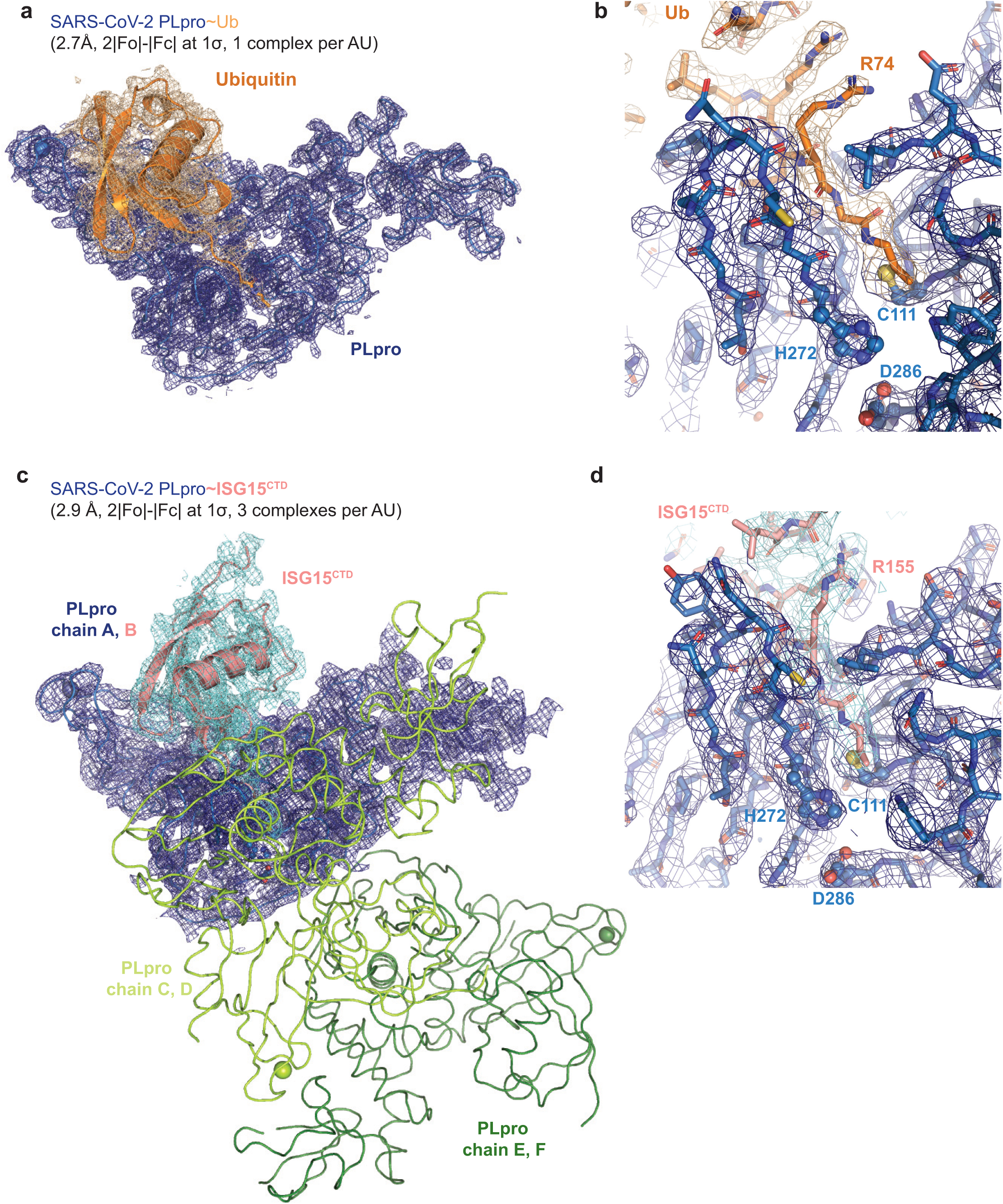
Electron density for PLpro complexes. **a**, 2|Fo|-|Fc| electron density, contoured at 1*σ*, for the PLpro∼Ub complex. The full asymmetric unit is shown. SARS2 PLpro is shown as a ribbon, and ubiquitin is shown in cartoon representation. **b**, Detailed electron density for the ubiquitin C-terminal tail in the active site, with the propargyl linked to catalytic Cys111. **c**, 2|Fo|-|Fc| electron density, contoured at 1*σ*, for the PLpro∼ISG15^CTD^ complex covering chain A (PLpro, ribbon) and B (ISG15^CTD^, cartoon). The remainder (chains C-F) of the asymmetric unit are shown as a ribbon without map. **d**, Detailed electron density for the ISG15 C-terminal tail in the active site, with the propargyl-linked to catalytic Cys111.

**Extended Data Figure 4.**
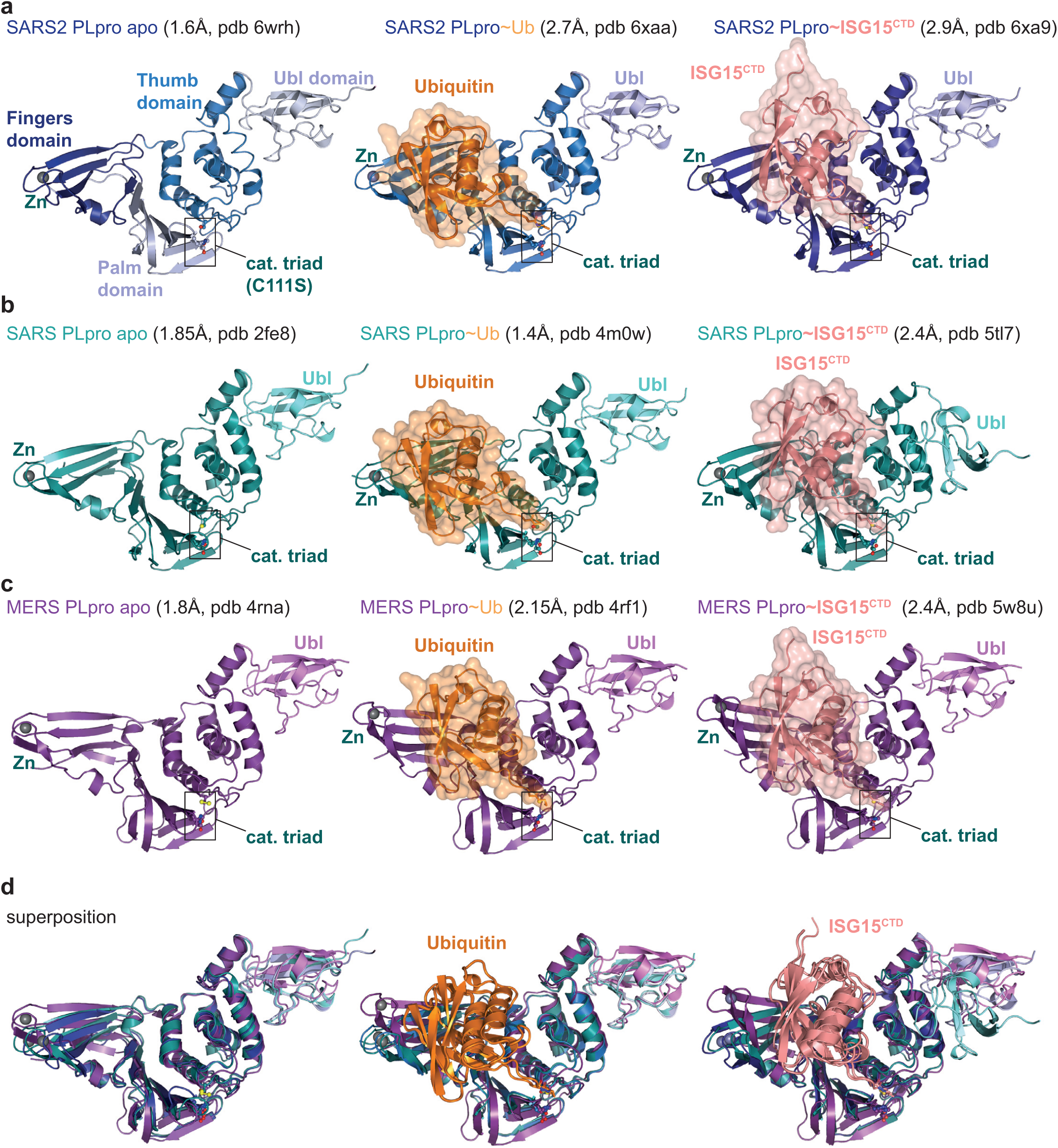
Selection of available structures of Coronavirus PLpro. A large body of work on SARS and MERS PLpro has led to determination of multiple structures of PLpro apo and PLpro bound to ubiquitin and ISG15. A selection of structures is displayed, when multiple structures were available, the highest resolution structure was used. **a**, *left*, the unpublished structure of apo SARS2 PLpro (pdb 6wrh, 1.6 Å, determined by the Centre for Structural Genomics of Infectious Disease (CSGID)) is coloured in analogy with **Figure 1d** and **Extended Data Fig. 1**, indicating Thumb, Fingers, and Palm subdomain. The PLpro fold forms an open right hand that holds the ubiquitin fold, guiding its C-terminus into the active site. PLpro contains an N-terminal Ubl domain of unknown function as the most structurally variable domain of PLpro. The high-resolution structure was generated with a catalytic Cys to Ser mutation. We found a more common Cys to Ala mutant of the catalytic Cys to result in highly unstable protein (**Extended Data Fig. 5d**). *Middle*, structure of SARS2 PLpro bound to ubiquitin (orange, covered by a semi-transparent surface). Also see **Figure 1**. *Right*, structure of SARS2 PLpro bound to the C-terminal domain of ISG15 (ISG15^CTD^, salmon, under a semi-transparent surface). Also see **Figure 1**. In ubiquitin and ISG15^CTD^ complexes, propargylamide based suicide probes ^41^ covalently modify catalytic Cys111. **b**, *left*, SARS PLpro apo (1.85 Å, pdb 2fe8, ref. ^56^), *middle*, SARS PLpro bound to ubiquitin (1.4 Å, pdb 4m0w, ref. ^23^), *right*, SARS PLpro bound to the C-terminal domain of ISG15 (2.4 Å, pdb 5tl7, ^25^). **c**, *left*, MERS PLpro apo (1.84 Å, pdb 4rna, ref. ^32^), *middle*, MERS PLpro bound to ubiquitin (2.15 Å, pdb 4rf1, ref. ^26^), *right*, MERS PLpro bound to the C-terminal domain of ISG15 (2.4 Å, pdb 5w8u, ref. ^57^). **d**, Superpositions of PLpro structures with S1 site occupied by different modifiers. Overall, PLpro shows high similarity, extending to the position and orientation of the N-terminal Ubl domain, with the notable exception of a distinct position of the Ubl in the SARS complex with ISG15^CTD^ (pdb 5tl7, ^25^). The second, most variable region concerns the Fingers subdomain, which shows varying degrees of ‘openness’. Superposition shows that the structures of SARS and SARS2 bound to individual modifiers are highly similar, and the modifiers adapt identical orientations and engage in similar interactions with PLpro. MERS PLpro seems to vary on the theme of ubiquitin versus ISG15 recognition, by binding both modifiers similarly. In the MERS ubiquitin complex, the fingers are more closed, and the ubiquitin is pushed towards the Thumb domain, to adopt a similar orientation and interaction as seen for ISG15 bound to MERS. MERS PLpro ubiquitin complexes have been determined with ‘open’ and more ‘closed’ Fingers ^26^.

**Extended Data Figure 5.**
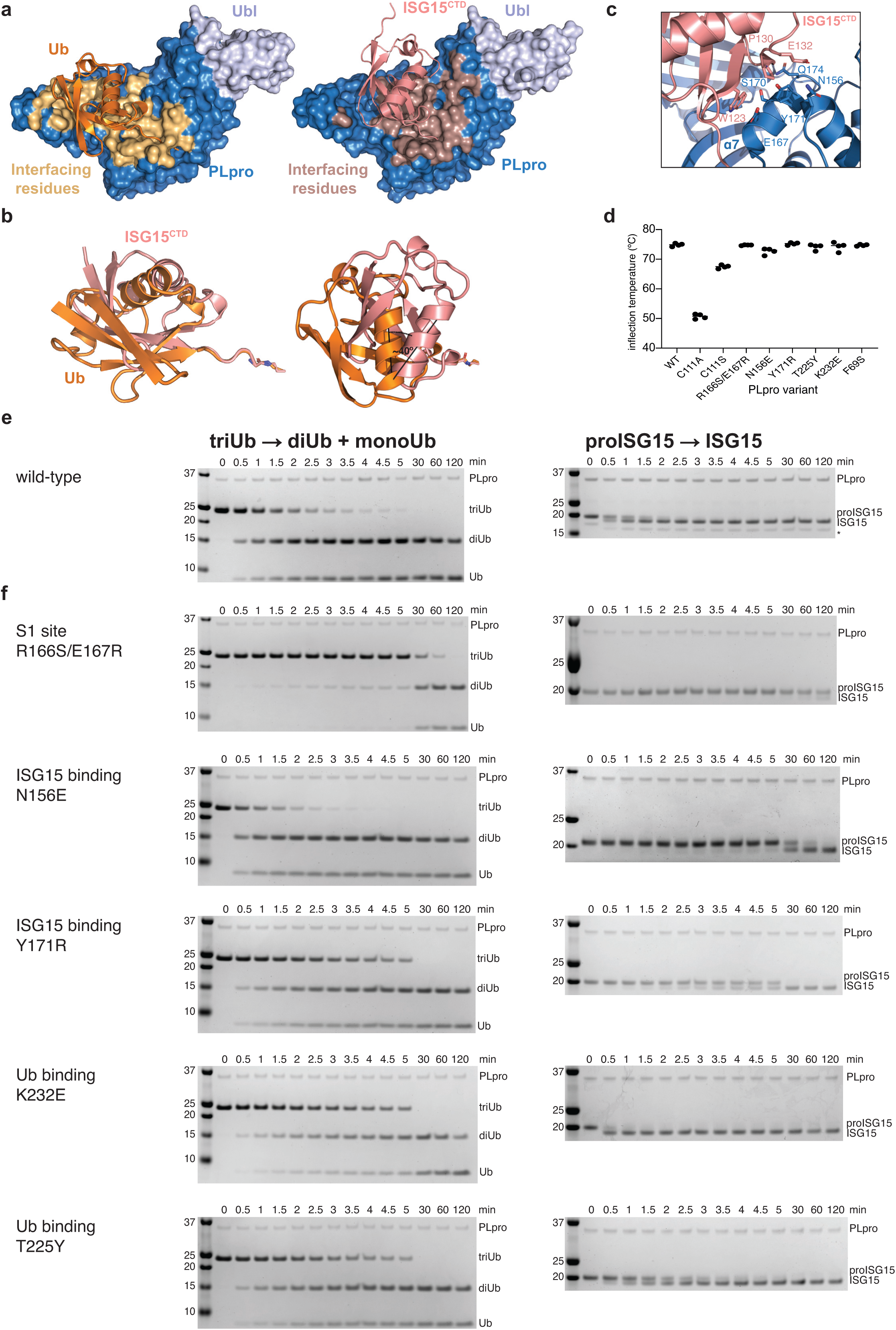
Separation of function mutations in SARS2 PLpro. **a**, Ubiquitin and ISG15 binding site analysis based on PISA analysis, indicating interface residues on SARS2 PLpro. **b**, Superposition of Ub-PA (orange) and ISG15^CTD^-PA (salmon) as bound to SARS2 PLpro highlights the different binding modes with a ∼40° rotation between the two proteins. **c**, Details of the binding of ISG15^CTD^ and the Thumb domain of SARS2 PLpro. Interacting residues shown as sticks. **d**, Mutations in S1 and S2 sites were introduced in PLpro, and the enzyme variants were expressed in bacteria, purified, and tested for integrity by assessing the inflection temperature, indicating the transition of folded to unfolded protein. With exception of mutating the catalytic Cys to Ala, which was severely destabilised and precipitated during purification, all other mutants showed similar stability to wild-type PLpro. Inflection temperature values were determined in technical duplicate from experiments performed twice. **e, f**, Triubiquitin cleavage to mono- and diubiquitin (*left*), and proISG15 cleavage to mature ISG15 (*right*), were compared side-by-side over a time course, resolved on SDS-PAGE gels, and visualised by Coomassie staining. Experiments were performed in duplicate with 250 nM enzyme and 2 µM substrate; all gels are shown in **Supplementary Figure 1. c**, Activity of wild-type PLpro, reproduced from **Extended Data Fig 2b, 2c**, for comparison. **d**, S1 site mutants as indicated. See **Figure 2**.

**Extended Data Figure 6.**
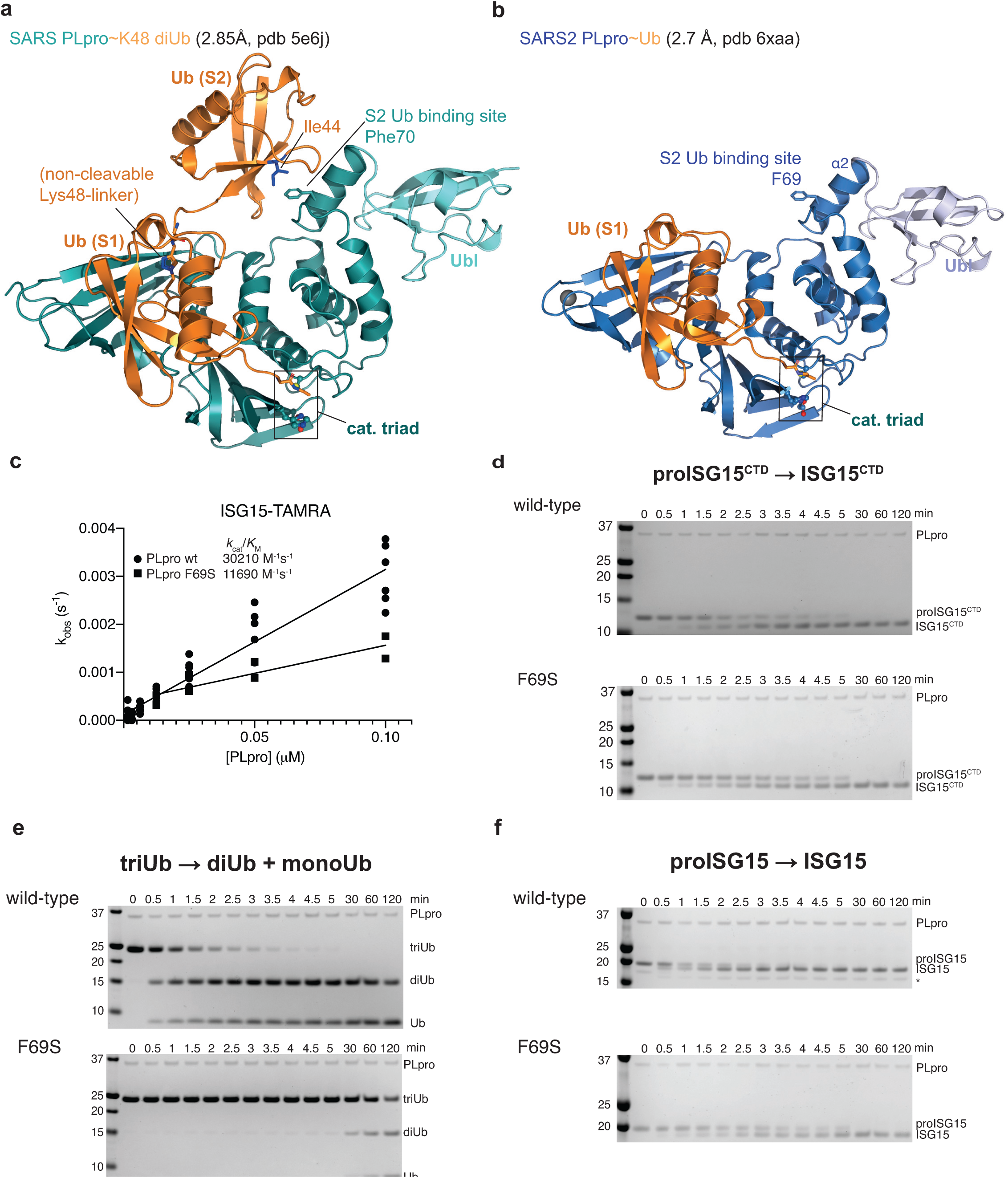
The S2 site in SARS2 Plpro. **a**, A previous structure of SARS PLpro bound to a non-hydrolysable, Lys48-linked diubiquitin probe (pdb 5e6j, ^20^) explained the noted preference of PLpro for longer Lys48-linked chains. While the proximal ubiquitin unit occupies the S1 site in a highly similar fashion in SARS∼Ub and SARS2∼Ub structures (see **b, Figure 2a, Extended Data Fig. 4**), the second, distal, ubiquitin unit binds to the α2 helix of PLpro, through a common binding mode involving the ubiquitin Ile44 patch. On helix α2, a central Phe70 in SARS PLpro residue is flanked by residues involved in polar contacts. **b**, Structure of the SARS2 PLpro∼Ub complex. The S2 site consisting of α2 helix with Phe69 residue, is fully conserved in SARS2 PLpro. Mutation of Phe69 to Ser severely impacts triubiquitin and proISG15 hydrolysis (see **Extended Data Fig. 5e**). **c**, Fluorescence polarisation assay on ISG15-TAMRA for PLpro wild-type (reproduced from **Extended Data Fig. 2a**) and PLpro F69S. A ∼3-fold lower efficiency for F69S is similar to cleavage of ISG15^CTD^-TAMRA (**Extended Data Fig. 2a**), suggesting that the S2 site contributes the difference in binding for the N-terminal Ubl-fold. Experiments for F69S were performed in technical triplicate and biological duplicate. **d-f**, Gel based analysis showing hydrolysis of proISG15^CTD^ (**d**), triubiquitin (**e**) and proISG15 (**f**) using wild-type PLpro (*top*, gels reproduced from **Extended Data Fig. 2b-d** to enable direct comparison) or PLpro F69S (*bottom*). Mutation of the S2 site has no marked effect on hydrolysis of proISG15^CTD^ (**d**) and reduces proISG15 cleavage to the same levels as proISG15^CTD^ (compare **f** and **d**), PLpro F69S has a dramatic effect on triubiquitin hydrolysis (**e**). Experiments were performed in duplicate, see **Supplementary Figure 1**.

**Extended Data Figure 7.**
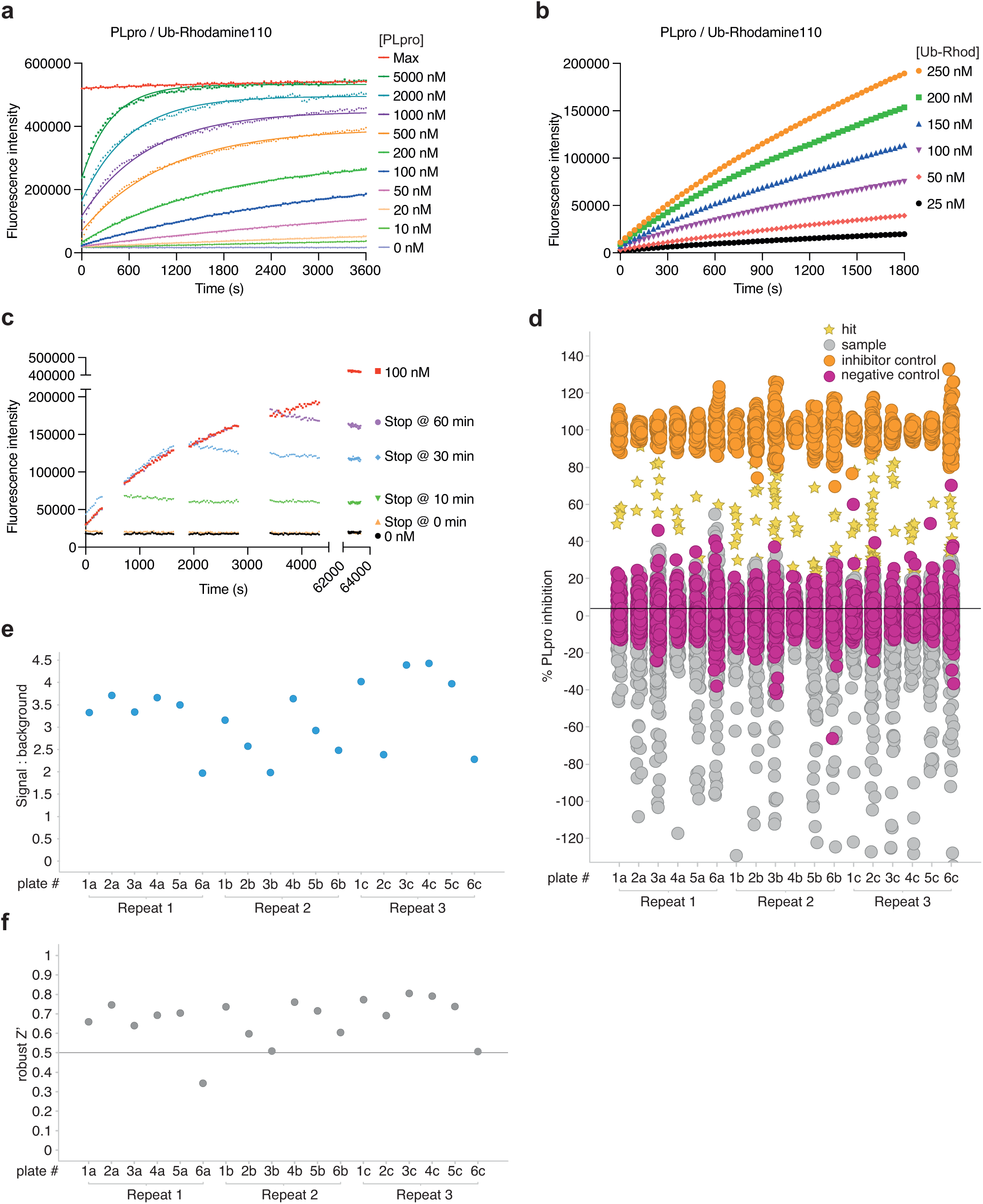
Developing a high-throughput screen to identify SARS2 PLpro inhibitors. **a**, Suitable SARS2 PLpro concentrations were determined by kinetic analysis of increasing Ub-Rhodamine fluorescence over 1 h (3600 s). Concentrations ranged from 10-5000 nM with sufficient signal obtained with 50 nM SARS2 PLpro at a constant concentration of 100 nM Ub-Rhodamine. Maximal signal (Max) indicates pre-incubated Ub-Rhodamine with 25 nM PLpro for 1 h, to achieve complete cleavage of Ub-Rhodamine, before measurement. **b**, To determine the optimal substrate concentration, 50 nM SARS2 PLpro was incubated with 25-250 nM Ub-Rhodamine for 30 min (1800 s). A final concentration of 100nM Ub-Rhodamine was selected, which was well below *K*_M_ for SARS2 PLpro, in the linear range of the reaction, with a signal to background (S:B) above 3 at 12 min (720 s). 12 min was the timepoint selected for end-point assays. **c**, Enzymatic reactions were stopped with addition of citric acid at a final concentration of 10 mM at indicated timepoints. The assay was benchmarked against compound rac5c (see below, **Figure 4** and **Extended Data Fig. 8**). Rac5c inhibited SARS2 PLpro activity with an IC_50_ of 0.81 µM (**Figure 4c**, see **Methods, Supplementary HTS Information**). **d**, Results from the complete screen, by plate number. **e**, Signal:background analysis from the whole screen by plate number. 17 out of 18 plates met the quality control criteria (S:B > 2). **f**, Robust Z’ analysis of the whole screen by plate number. Plate 6a, which did not meet quality criteria in S:B and robust Z’ analysis, was excluded from analysis.

**Extended Data Figure 8.**
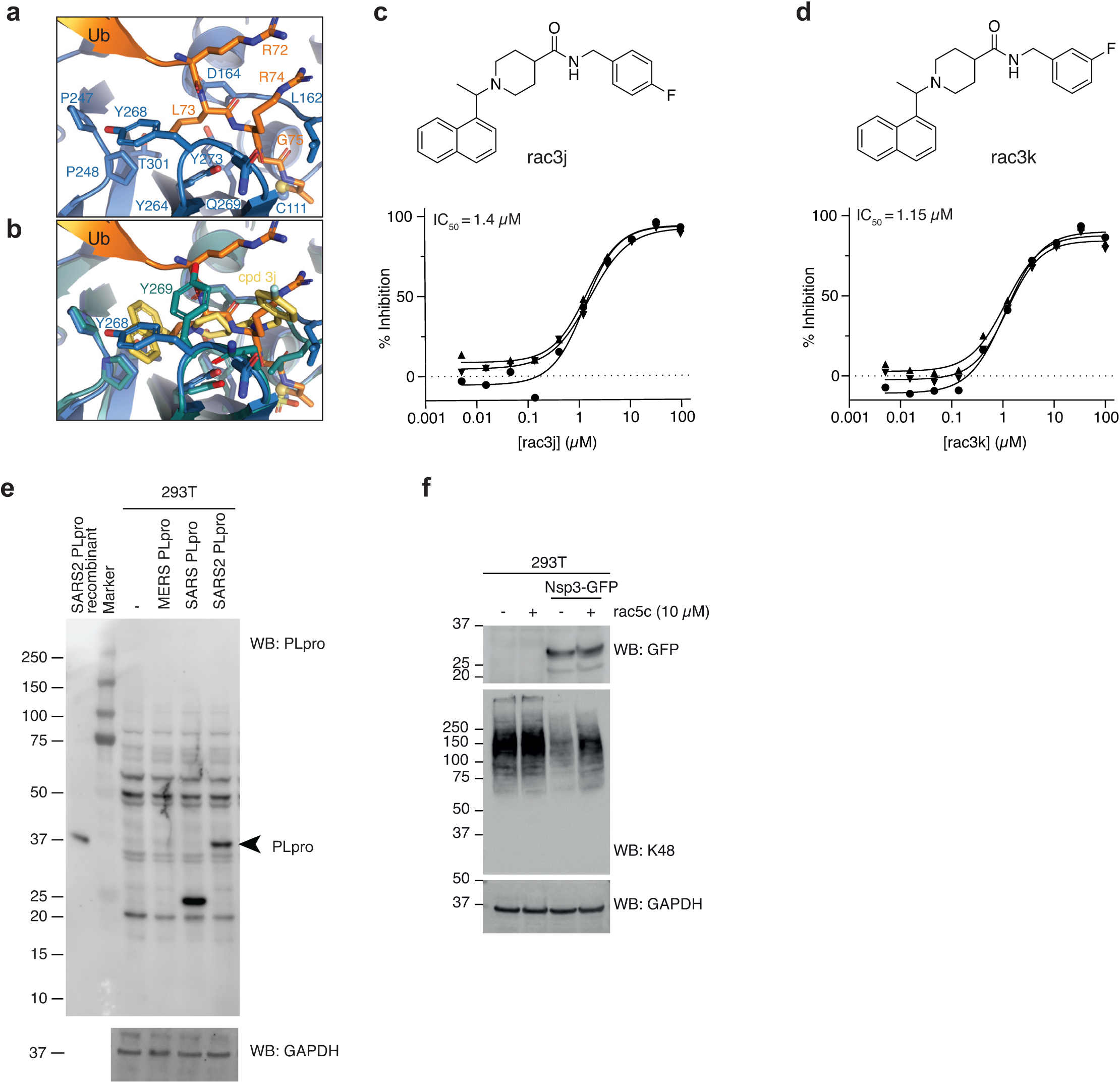
SARS PLpro compounds inhibit SARS2 Plpro. **a**, Structure of SARS2 PLpro bound to the ubiquitin C-terminal tail in the active site, compare with **Figure 4a. b**, Superposition of ubiquitin tail in SARS2 PLpro, and compound 3j in SARS PLpro (pdb 4ovz, ^29^) shows an identical binding for compounds in SARS2 PLpro and highlights the change in Tyr268/269 in SARS2 PLpro and SARS PLpro, respectively. **c, d**, Compounds rac3j and rac3k, racemic versions of 3j and 3k from ^29^, and their *in vitro* biochemical IC_50_ values determined by the HTS assay technical triplicate and in three independent experiments (as for rac5c in **Figure 4c**). **e**, Immunoblot characterisation of the PLpro antibody on HEK 293T cells overexpressing PLpro from MERS, SARS or SARS2. Cell lysates were immunoblotted 48 h post transfection. **f**, Immunoblot analysis showing the effect of rac5c (10 µM for 24 h) on Lys48-polyubiquitin chain disassembly by nsp3, 48 h post transfection in HEK 293T cells. Note that compounds have no effect on Lys48 chains in untransfected HEK293T cells.

**Extended Data Figure 9.**
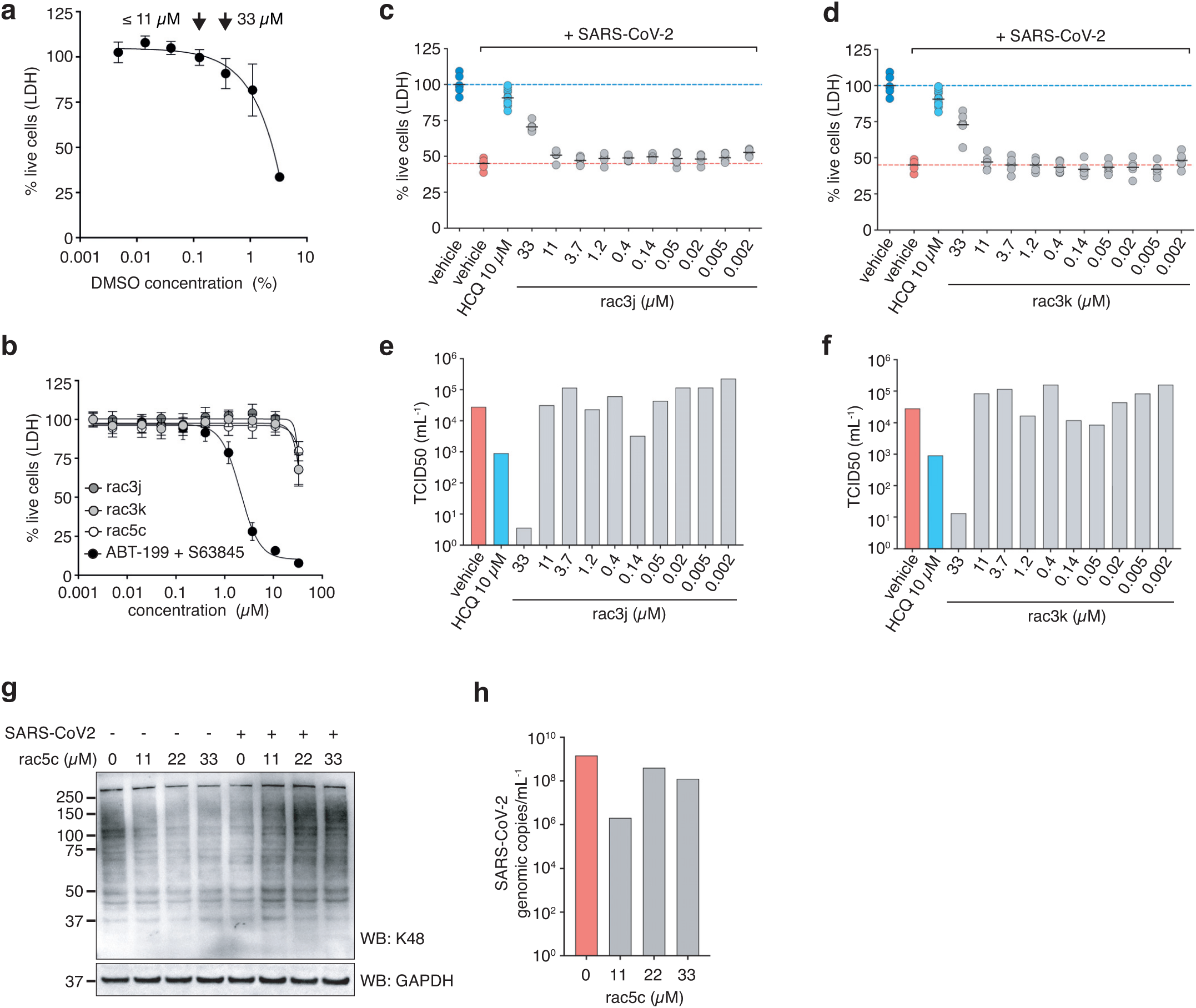
Antiviral activity of compounds in cells. **a**, Vero cells were tested for compatibility with DMSO concentrations, revealing low toxicity at concentrations <0.1% (v/v) but more substantial cytotoxicity at higher concentrations (0.3% and above). This limited the range of compound concentrations useable for infection studies; at concentrations up to 11 µM, 0.1% (v/v) DMSO was used as vehicle, at 33 µM compound concentration, 0.3% (v/v) DMSO was used. Higher concentrations of compound could not be tested due to this limitation. **b**, Toxicity titration of compounds rac3j, rac3k, and rac5c on Vero cells. At 33 µM compound concentration, cellular toxicity is ∼25%. ABT-199 and S63845 are death-inducing compounds ^58,59^ used as a control. **c, d**, CPE assays to assess cell killing activity of SARS-CoV-2 in Vero cells left untreated (DMSO control), or treated with compounds rac3j (**c**) and rac3k (**d**). One experiment with 6 biological replicates is shown (black line, mean), and compared to HCQ treated cells (10 µM) with pooled data from 2 experiments with n=6, as in **Figure 5b**. The 33 µM compound concentration was performed at 0.3% (v/v) DMSO and significantly rescued infected cells despite underlying cytotoxicity in Vero cells (see **a,b**). **e, f**, TCID50 analysis of infectious virus for rac3j, rac3k, from the experiments performed in **c** and **d**, respectively. Data is representative of 1 experiment out of 2, with n=6 per experiment, bars represent the mean TCID50 value. HCQ control was performed within the same experiment. **g**, Calu3 cells were infected with SARS-CoV-2 and treated 20 h post infection with increasing concentrations of rac5c for 4 h. Total cell lysates were blotted for Lys48-polyubiquitin with a linkage specific antibody and reprobed with for GAPDH as loading control. See **Supplementary Figure 1** for uncropped blots. **h**, SARS-CoV-2 infected and rac5c treated Calu-3 cells were sampled before lysis for Western blot analysis (for **g**) and RNA was extracted from total cell lysate. Reverse transcribed cDNA was analysed for virus specific RNA by qRTPCR (see **Methods**).

**Extended Data Table 1.** Data collection and refinement statistics. Values in parentheses are for highest-resolution shell.

|  | SARS2 PLpro~Ub | SARS2 PLpro~ISG15<sup>CTD</sup> |
| --- | --- | --- |
| **Data collection** |  |  |
| Space group | *P* 2<sub>1</sub> 2<sub>1</sub> 2 | *P* 4<sub>1</sub> 2<sub>1</sub> 2 |
| Cell dimensions |  |  |
| *a*, *b*, *c* (Å) | 64.99, 144.41, 49.60 | 124.17, 124.17, 238.17 |
| *α*, *β*, *γ* (°) | 90.00, 90.00, 90.00 | 90.00, 90.00, 90.00 |
| Resolution (Å) | 48.30 – 2.70 (2.83 – 2.70) | 49.28 – 2.90 (3.01 – 2.90) |
| *R*<sub>merge</sub> (within I+/I-) | 0.152 (1.054) | 0.163 (2.876) |
| < I/σI > | 6.8 (1.8) | 10.5 (1.1) |
| Completeness (%) | 98.1 (98.9) | 100.0 (100.0) |
| Redundancy | 4.6 (4.7) | 13.7 (14.2) |
| **Refinement** |  |  |
| Resolution (Å) | 48.30 – 2.70 | 49.28 – 2.90 |
| No. reflections | 13004 | 42059 |
| *R*<sub>work</sub> / *R*<sub>free</sub> | 0.210/ 0.260 | 0.200/ 0.231 |
| No. atoms |  |  |
| Protein | 2986 | 8876 |
| Ligand/ion | 5 | 63 |
| Water | 32 | 30 |
| *B*-factors |  |  |
| Protein | 58.2 | 102.1 |
| Ligand/ion | 53.1 | 101.7 |
| Water | 45.8 | 77.9 |
| R.m.s. deviations |  |  |
| Bond lengths (Å) | 0.0024 | 0.0059 |
| Bond angles (°) | 0.66 | 0.94 |

All data sets were collected from a single crystal each. Values in parentheses are for highest-resolution shell.

