## Supplementary material for "Mechanism and inhibition of SARS-CoV-2 PLpro": Screening Table

| Category | Parameter | Description |
| --- | --- | --- |
| Assay | Type of assay | Enzymatic, endpoint |
| | Target | SARS-CoV-2 Papain-like Protease |
| | Primary measurement | Fluorescence intensity |
| | Key reagents | Ub-Rhodamine |
| Library | Library size | 5,576 compounds |
| | Library composition | Small molecule, including FDA-approved drugs, advanced preclinical compounds and focused sets. |
| | Source | Commercial and in-house curated collections (see Methods) |
| | Additional comments | |
| Screen | Format | 1536 well plate |
| | Concentration tested | 4.2 μM in 2% DMSO |
| | Plate controls | Negative control; 2% DMSO, positive control; compound rac5c (see manuscript, Figure 4) |
| | Reagent/ compound dispensing system | See Methods |
| | Detection instrument and software | PHERAstar FSX (BMG) |
| | Assay validation/QC | Robust Z' > 0.5 |
| | Hit criteria | >4*MAD above negative control |
| | Hit rate | 0.45% |
| Post-HTS analysis | Counter screen | Human USP21, Ub-Rhodamine |
| | Additional assay(s) | |
| | Confirmation of hit purity and structure | LC/MS |
