## Supplementary material for "Mechanism and inhibition of SARS-CoV-2 PLpro": Uncropped Gels

Supplementary Information Figure 1

#### Uncropped versions of SDS-PAGE gels

PLpro WT Ub chain specificity assays

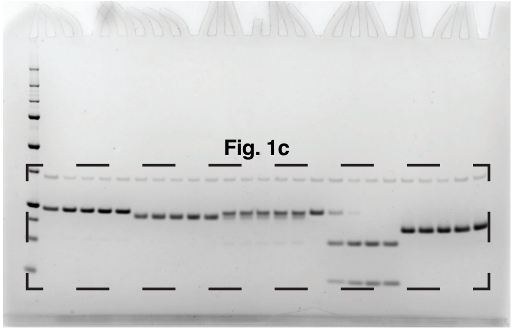


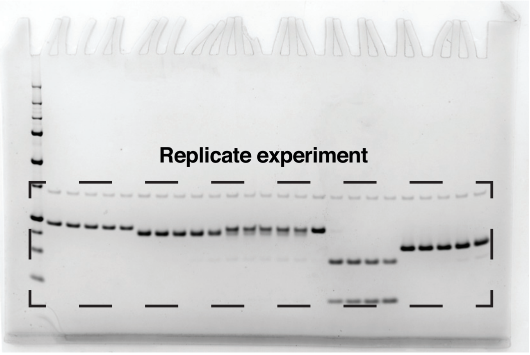


S1 site WT/Mts + K48 triUb kinetics comparison

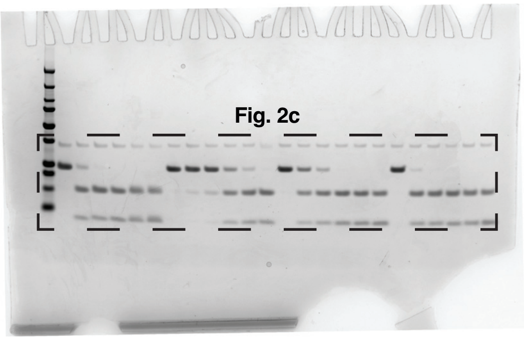


S1 site WT/Mts + proISG15 kinetics comparison

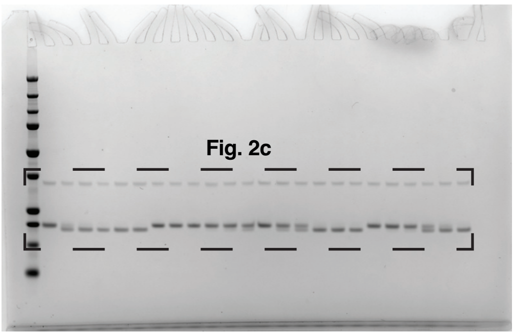


PLpro WT + K48 triUb kinetics assays

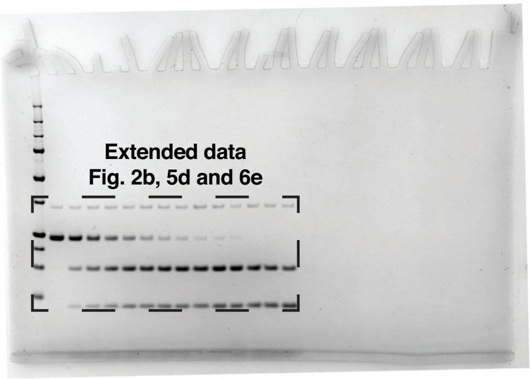


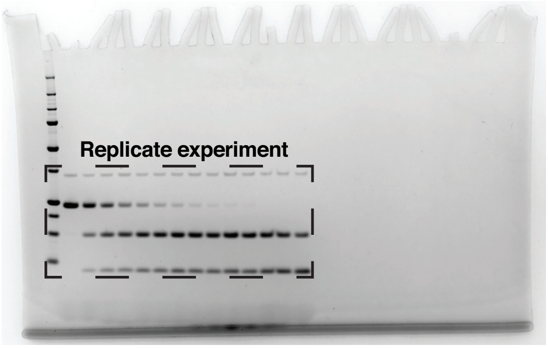


PLpro WT + proISG15 kinetics assays

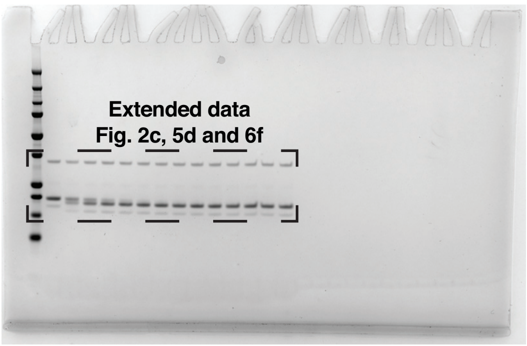


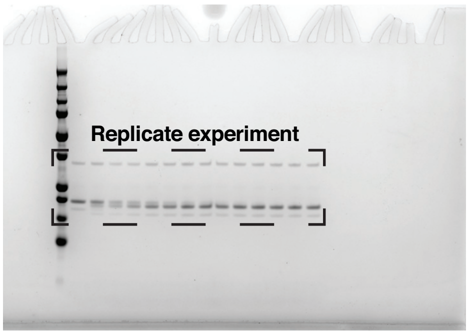


PLpro R166S/E167R + K48 triUb kinetics assays

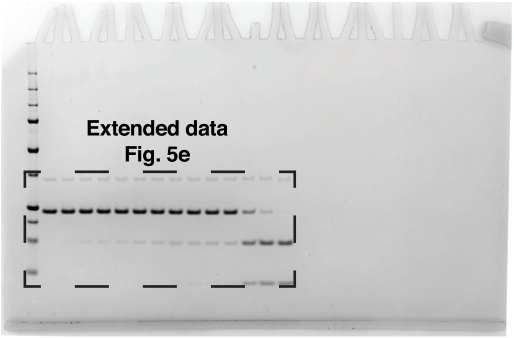


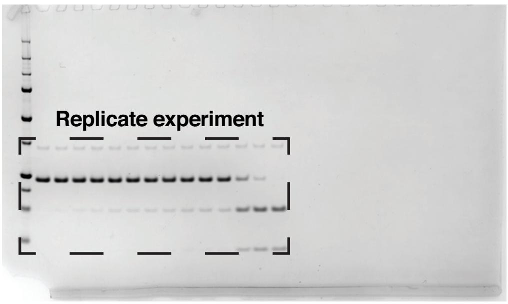

### Supplementary Information Figure 1

PLpro R166S/E167R + proISG15 kinetics assays

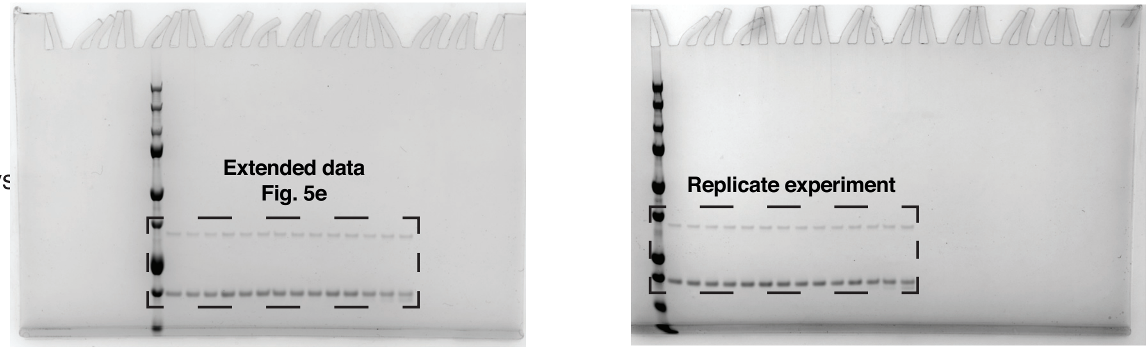


PLpro N156E + K48 triUb kinetics assays

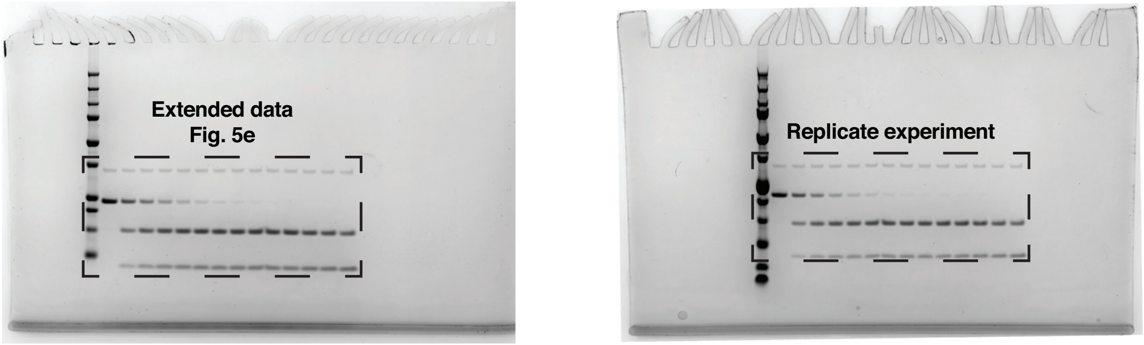


PLpro N156E + proISG15 kinetics assays

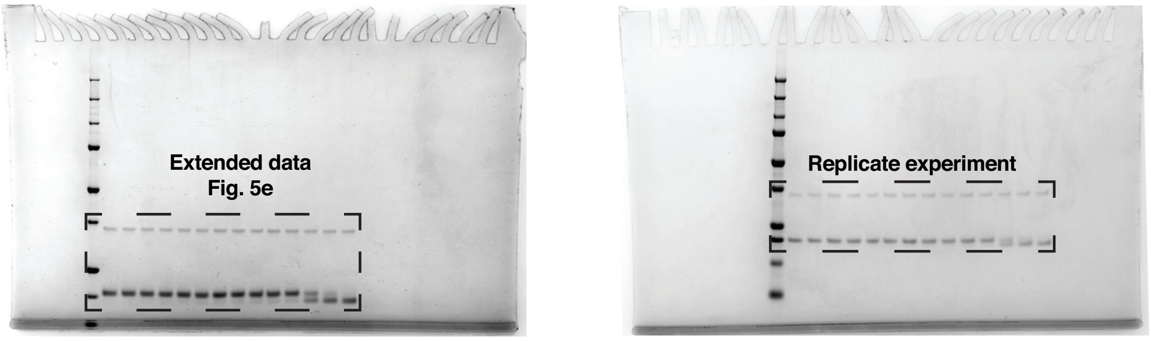


PLpro Y171R + K48 triUb kinetics assays

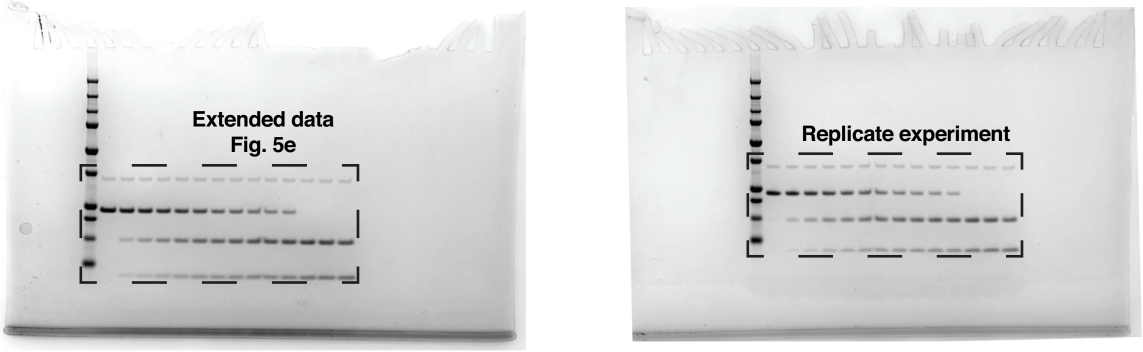


PLpro Y171R + proISG15 kinetics assays

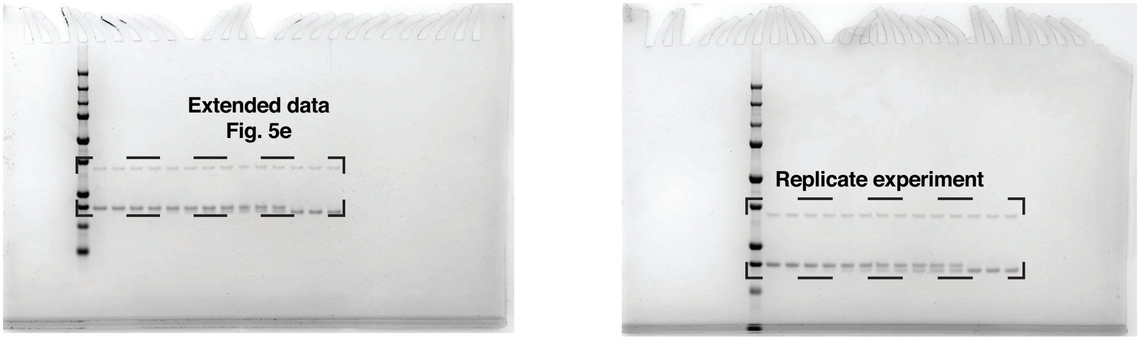

### Supplementary Information Figure 1

PLpro K232E + K48 triUb kinetics assay

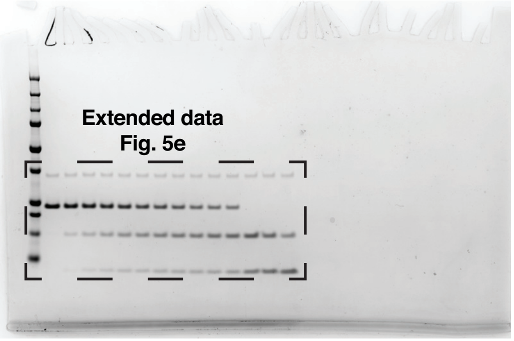


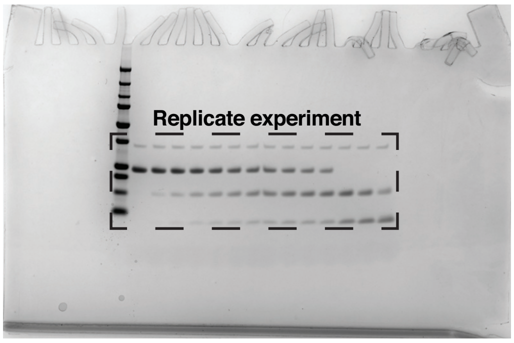


PLpro K232E + proISG15 kinetics assays

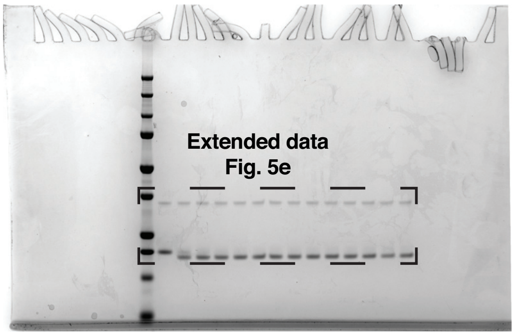


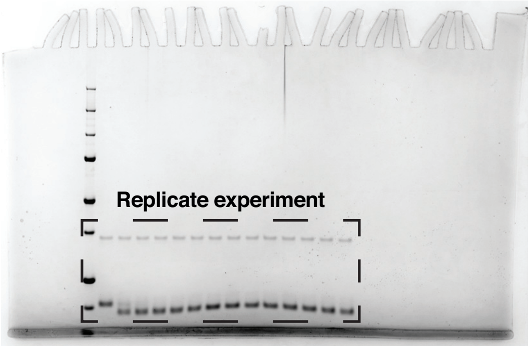


PLpro T225Y + K48 triUb kinetics assays

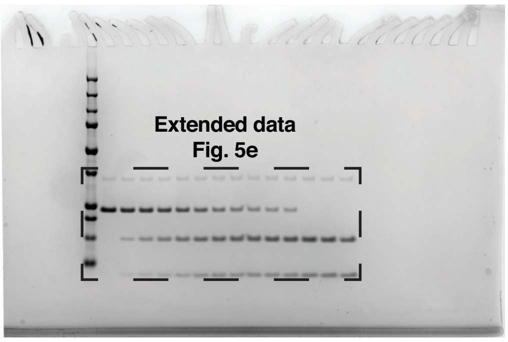


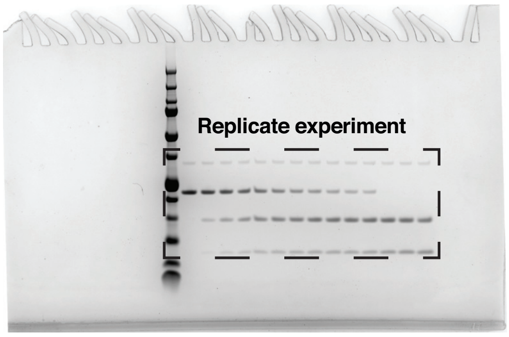


PLpro T225Y + proISG15 kinetics assays

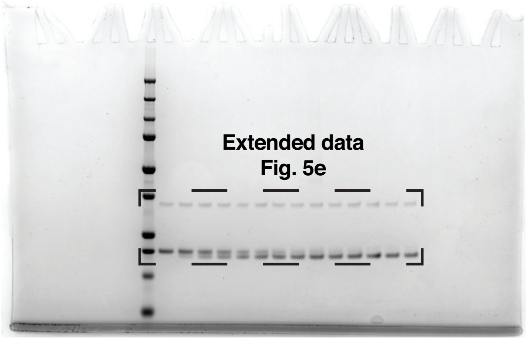


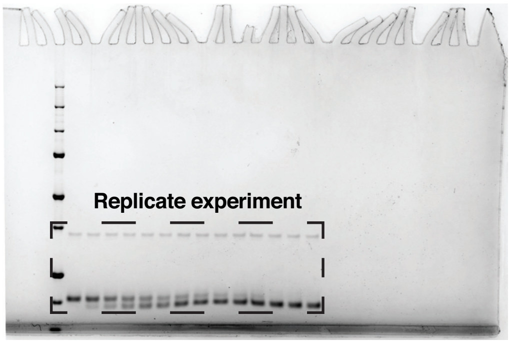


PLpro F69S + K48 triUb kinetics assays

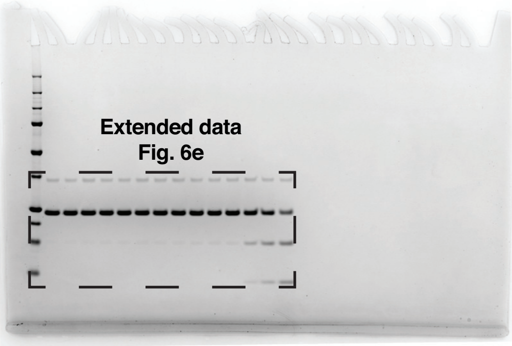


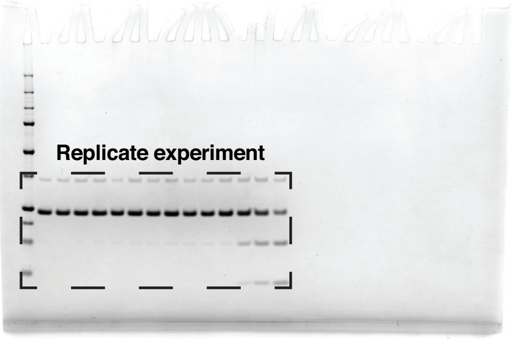

PLpro F69S + proISG15 kinetics assays

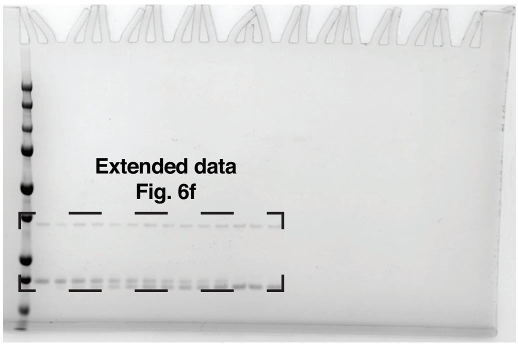


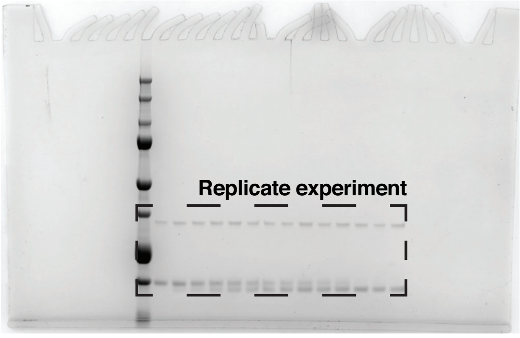


PLpro WT + ISG15$^{CTD}$ kinetics assays

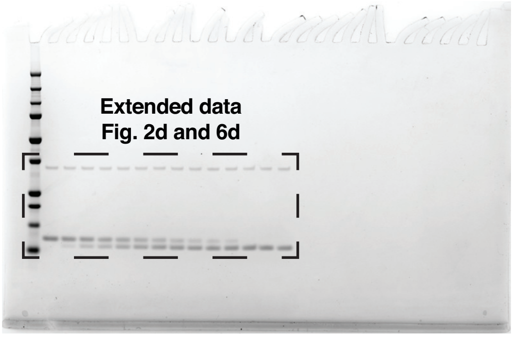


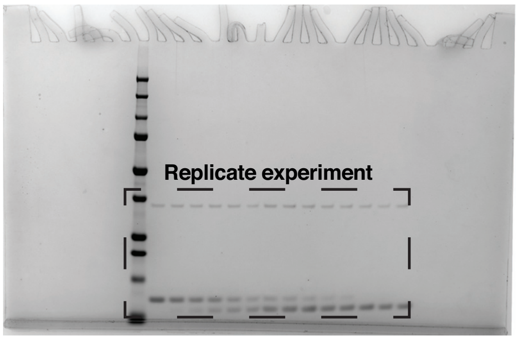


PLpro F69S + proISG15$^{CTD}$ kinetics assays

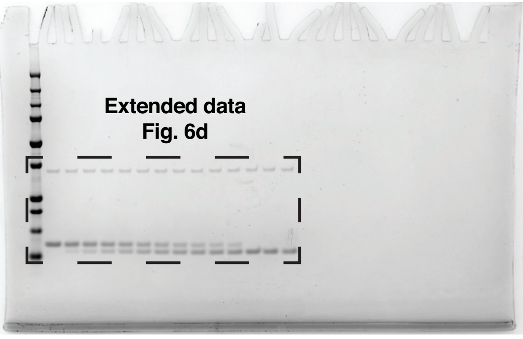


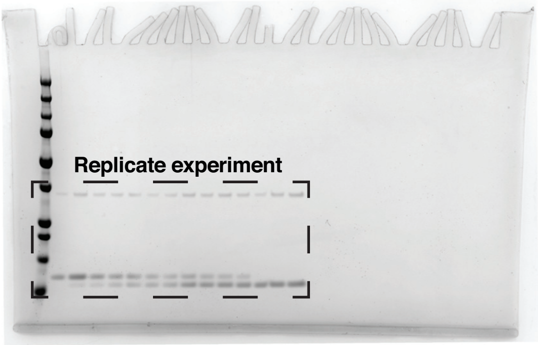

Supplementary Information Figure 1

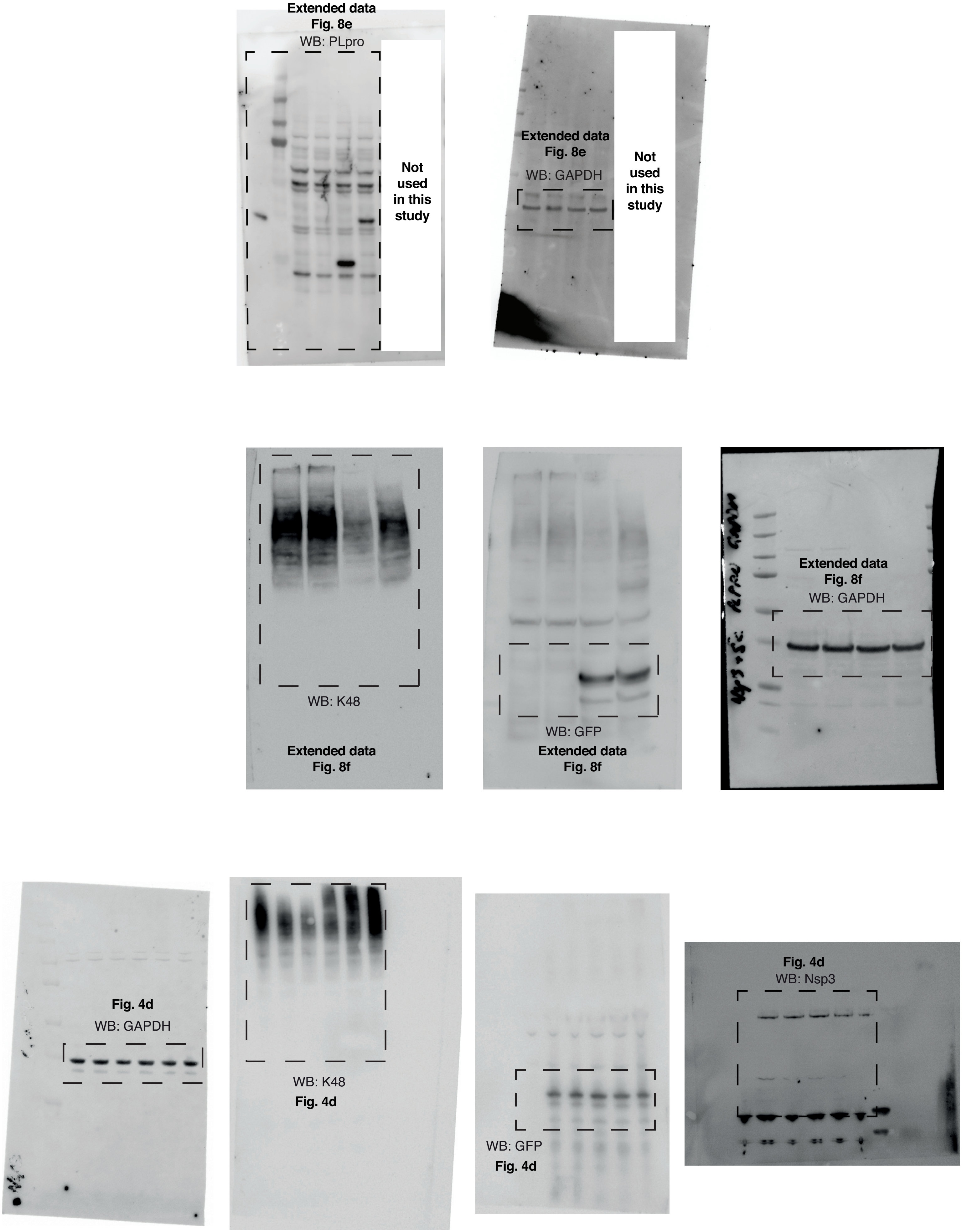

Supplementary Information Figure 1

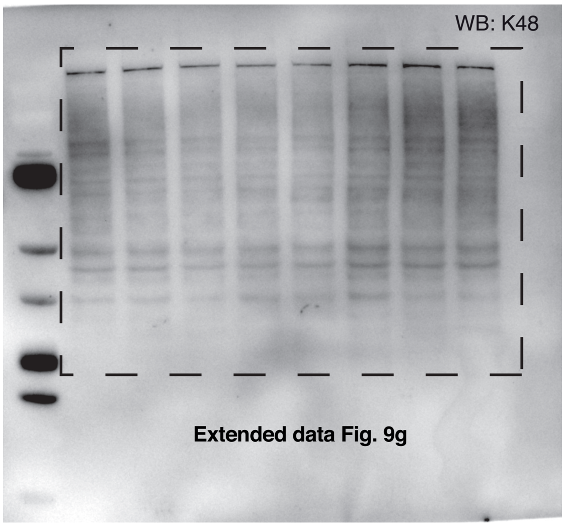


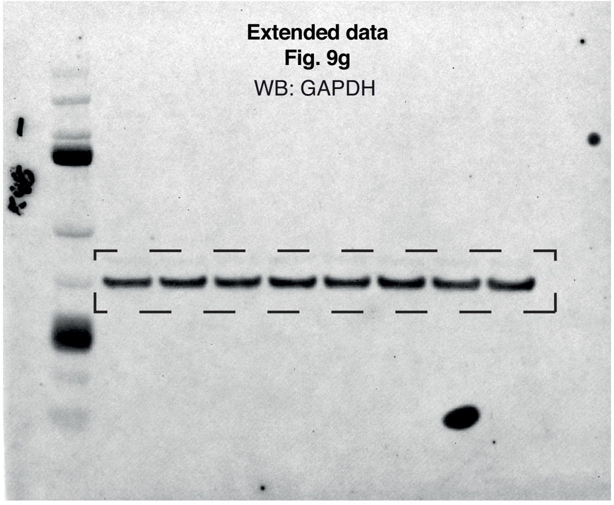
