## Supplementary material for "Mechanism and inhibition of SARS-CoV-2 PLpro": Chemistry Methods

**Supplemental Experimental Procedures**

**General chemistry**

Anhydrous solvents were obtained commercially (SIGMA-ALDRICH, Missouri, USA) and used without further purification. All other commercial reagents were used as supplied. All non-aqueous reactions were performed in oven-dried glassware under inert atmosphere (nitrogen gas), unless otherwise specified. Analytical thin-layer chromatography was performed on silica gel $^{60}F_{254}$ aluminum-backed plates (MERCK MILLIPORE) and were visualised by fluorescence quenching under UV light or by $KMnO_4$ staining. Chromatography was performed with silica gel 60 (particle size 0.040 – 0.063 µm) using an automated purification system (ISCO TELEDYNE). NMR spectra were recorded on a Bruker Ascend-300 300 MHz at 298 K unless otherwise specified. Chemical shifts are reported in ppm on the δ scale and referenced to the appropriate solvent peak. DMSO-$d_6$, MeOD and $CDCl_3$ contain $H_2O$. HRMS analyses were carried out at the Monash University Mass Spectrometry Facility on an Agilent 6224 TOF LC/MS Mass Spectrometer coupled to an Agilent 1290 Infinity (Agilent, Palo Alto, CA). All data were acquired and reference mass corrected via a dual-spray electrospray ionisation (ESI) source. LCMS were recorded on an Agilent LCMS system composed of an Agilent G6120B Mass Detector, 1260 Infinity G1312B Binary pump, 1260 Infinity G1367E HiPALS autosampler and 1260 Infinity G4212B Diode Array Detector. Conditions for LCMS were as follows, (Method A) column: Poroshell 120 EC-C18, 2.1 x 50 mm 2.7 Micron at 20 °C, injection volume 2 µL, gradient: 5–100% B over 3 min (solvent A: water 0.1% formic acid; solvent B: acetonitrile 0.1% formic acid), flow rate: 0.8 mL/min, detection: 254 nm, acquisition time: 5 min. HPLC conditions used to assess purity of final compounds were as follows, column: Phenomenex Gemini C18, 2.0 x 50 mm; injection volume 20 µL; gradient: 0–100% Buffer B over 6 min (buffer A: 0.1% formic acid in autoclaved MilliQ water; buffer B: 0.1% formic acid in 100% acetonitrile), flow rate: 1.0 mL/min, detection: 214 or 224 nm.

**Experimental – Racemic compounds**

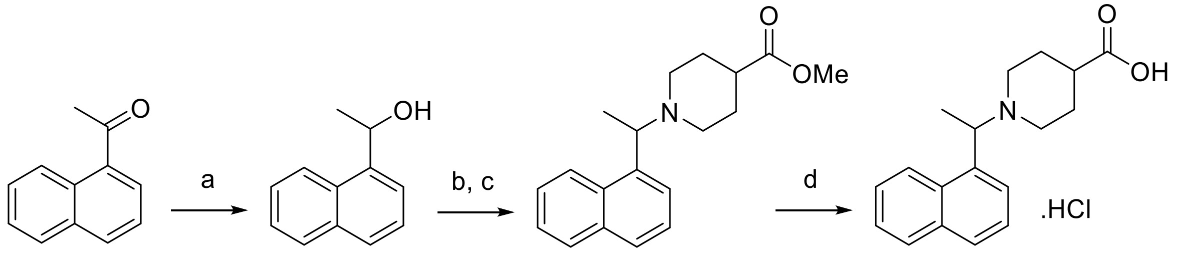

1 2 3 4

**Step (a)**

To a 0 °C solution of 1'-acetonaphthone (1.1 g, 6.5 mmol) in EtOH (20 mL) was added $NaBH_4$ (286 mg, 7.6 mmol) in one portion. The cooling bath was removed, and the reaction stirred until complete by LCMS. The reaction was concentrated under reduced pressure and the crude partitioned between saturated aqueous $NH_4Cl$ and EtOAc. The layers were separated and the aqueous further extracted with EtOAc. The combined organics were dried over anhydrous $MgSO_4$, filtered and concentrated under reduced pressure. The crude residue was subject to purification by combi-flash chromatography (0-40% EtOAc/n-heptane) to give 1-(naphthalen-1-yl)ethanol as a white solid (1.1 g, 98%).

1H NMR (300 MHz, CDCl3) δ 8.17 – 8.08 (m, 1H), 7.94 – 7.82 (m, 1H), 7.79 (dq, J = 8.2, 0.8 Hz, 1H), 7.69 (dt, J = 7.2, 1.0 Hz, 1H), 7.59 – 7.43 (m, 3H), 5.69 (q, J = 6.7 Hz, 1H), 1.98 (s, 1H), 1.68 (d, J = 6.4, 0.4 Hz, 3H).

**Steps (b, c)**

To a 0 °C solution of 1-(naphthalen-1-yl)ethanol (1.1 g, 6.4 mmol) in dry $CH_2Cl_2$ (20 mL) was added $SOCl_2$ (500 µL, 6.9 mmol) drop wise followed by catalytic DMF (1 drop). The reaction was stirred until complete by TLC. The reaction mixture was quenched with saturated aqueous $NH_4Cl$ and extracted with $CH_2Cl_2$. The combined organics were dried over anhydrous $MgSO_4$, filtered and concentrated under reduced pressure to give the desired alkyl chloride without any further purification. The crude alkyl chloride was taken up in dry MeCN (10 mL) and sequentially was added $K_2CO_3$ (1.5 g, 10.8 mmol) and methyl isonipectoate (960 µL, 7.1 mmol). The reaction was stirred at 70 ºC until complete by LCMS. The reaction was concentrated under reduced pressure and the crude residue partitioned between saturated aqueous $NH_4Cl$ and EtOAc. The layers were separated and the aqueous further extracted with EtOAc. The combined organics were dried over anhydrous $MgSO_4$, filtered and concentrated under reduced pressure. The crude residue was subject to purification by combi-flash chromatography (0-25% EtOAc/$CH_2Cl_2$) to give methyl 1-(1-(naphthalen-1-yl)ethyl)piperidine-4-carboxylate as a yellow oil (574 mg, 37%).

1H NMR (300 MHz, $CDCl_3$) δ 8.48 – 8.39 (m, 1H), 7.90 – 7.79 (m, 1H), 7.74 (d, J = 8.0 Hz, 1H), 7.57 (d, J = 7.1 Hz, 1H), 7.54 – 7.37 (m, 3H), 4.10 (q, J = 6.7 Hz, 1H), 3.66 (s, 3H), 3.19 – 3.09 (m, 1H), 2.87 – 2.78 (m, 1H), 2.37 – 2.21 (m, 1H), 2.16 – 1.98 (m, 2H), 1.97 – 1.87 (m, 1H), 1.85 – 1.66 (m, 3H), 1.46 (d, J = 6.7 Hz, 3H).

**Step (d)**

To a stirred solution of methyl 1-(1-(naphthalen-1-yl)ethyl)piperidine-4-carboxylate (564mg, 1.9mmol) in THF/MeOH (3:2) was added 40% aqueous NaOH (1 mL). The reaction was warmed at 60 °C until complete by LCMS. The reaction was concentrated under reduced pressure and the pH adjusted to 2 with 5N HCl. The precipitated solids were collected by filtration, washed with chilled $H_2O$ and dried to give 1-(1-(naphthalen-1-yl)ethyl)piperidine-4-carboxylic acid hydrochloride as a white solid (552 mg, 91%).

1H NMR (300 MHz, MeOD) δ 8.30 (d, J = 8.6 Hz, 1H), 8.08 – 7.96 (m, 2H), 7.91 – 7.83 (m, 1H), 7.75 – 7.55 (m, 3H), 5.44 (d, J = 6.4 Hz, 1H), 4.03 (s, 1H), 3.29 – 3.09 (m, 2H), 2.98 (s, 1H), 2.59 (s, 1H), 2.43 – 1.62 (m, 8H).

**N-(3-fluorobenzyl)-1-(1-(naphthalen-1-yl)ethyl)piperidine-4-carboxamide (rac3k)**

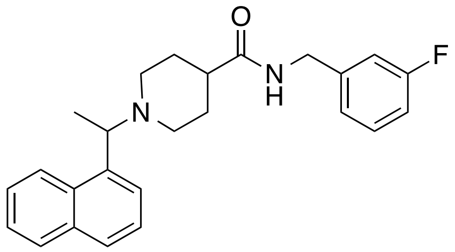

**rac3k**

To a solution of **4** (54 mg, 0.17 mmol) in dry DMF was added sequentially DIEA (62 mL, 0.36 mmol), ((3-fluorophenyl)methanamine (21 mL, 0.18 mmol) and HATU (71mg, 0.19mmol). The reaction was stirred at ambient temperature until complete by LCMS. The reaction was diluted with saturated aqueous $NaHCO_3$ and extracted with EtOAc. The combined extracts were washed with $H_2O$ (x3) and brine, dried over anhydrous $MgSO_4$, filtered and concentrated under reduced pressure. The crude residue was subject to purification by combi-flash chromatography (0-5% MeOH/$CH_2Cl_2$) to give the title compound (59 mg, 90%).

1H NMR (300 MHz, $CDCl_3$) δ 8.48 – 8.39 (m, 1H), 7.90 – 7.79 (m, 1H), 7.74 (d, *J* = 8.1 Hz, 1H), 7.56 (d, *J* = 7.1 Hz, 1H), 7.53 – 7.37 (m, 3H), 7.34 – 7.21 (m, 1H), 7.06 – 6.89 (m, 3H), 5.76 (s, 1H), 4.43 (d, *J* = 5.8 Hz, 2H), 4.11 (q, 1H), 3.24 (d, *J* = 11.3 Hz, 1H), 2.90 (d, *J* = 11.5 Hz, 1H), 2.26 – 1.65 (m, 7H), 1.47 (d, *J* = 6.7 Hz, 3H). ES+ MS: (M + H) 391.2. HPLC $t_g$ = 1.5 min, >95% purity.

**N-(4-fluorobenzyl)-1-(1-(naphthalen-1-yl)ethyl)piperidine-4-carboxamide (rac3j)**

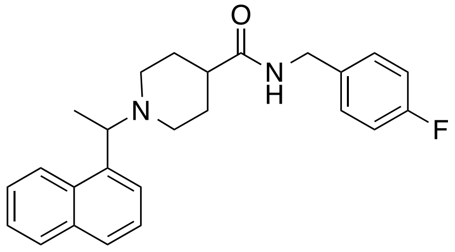

**rac3j**

The title compound **rac3j** was obtained as described for compound **rac3k** in 84% yield. $^1$H NMR (300 MHz, $CDCl_3$) δ 8.42 (d, $J$ = 7.8 Hz, 1H), 7.89 – 7.79 (m, 1H), 7.74 (d, $J$ = 8.1 Hz, 1H), 7.56 (d, $J$ = 7.1 Hz, 1H), 7.52 – 7.37 (m, 3H), 7.27 – 7.15 (m, 2H), 7.06 – 6.93 (m, 2H), 5.72 (s, 1H), 4.43 – 4.35 (m, 2H), 4.14 – 4.04 (m, 1H), 3.23 (d, $J$ = 11.1 Hz, 1H), 2.89 (d, $J$ = 11.4 Hz, 1H), 2.21 – 1.65 (m, 7H), 1.46 (d, $J$ = 6.7 Hz, 3H). ES+ MS: (M + H) 391.2. HPLC $t_g$ = 1.5 min, >95% purity.

**N-((2-methoxypyridin-4-yl)methyl)-1-(1-(naphthalen-1-yl)ethyl)piperidine-4-carboxamide (rac5c)**

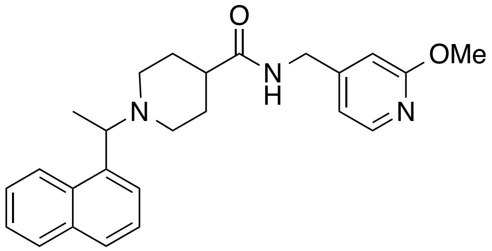


**rac5c**

The title compound **rac5c** was obtained as described for compound **rac3k** in 74% yield. $^1$H NMR (300 MHz, $CDCl_3$) δ 8.41 (br s, 1H), 8.08 (d, $J$ = 5.3 Hz, 1H), 7.90 – 7.81 (m, 1H), 7.75 (d, $J$ = 8.2 Hz, 1H), 7.58 (s, 1H), 7.54 – 7.38 (m, 3H), 6.77 – 6.69 (m, 1H), 6.58 (s, 1H), 5.84 (s, 1H), 4.39 (d, $J$ = 6.0 Hz, 2H), 4.15 (s, 1H), 3.91 (s, 3H), 3.31 – 3.21 (m, 1H), 2.98 – 2.88 (m, 1H), 2.22 – 1.71 (m, 7H), 1.55 – 1.43 (m, 3H). ES+ MS: (M + H) 404.2. HPLC $t_g$ = 1.2 min, >95% purity.

### NMR spectra of key compounds

#### Rac3j

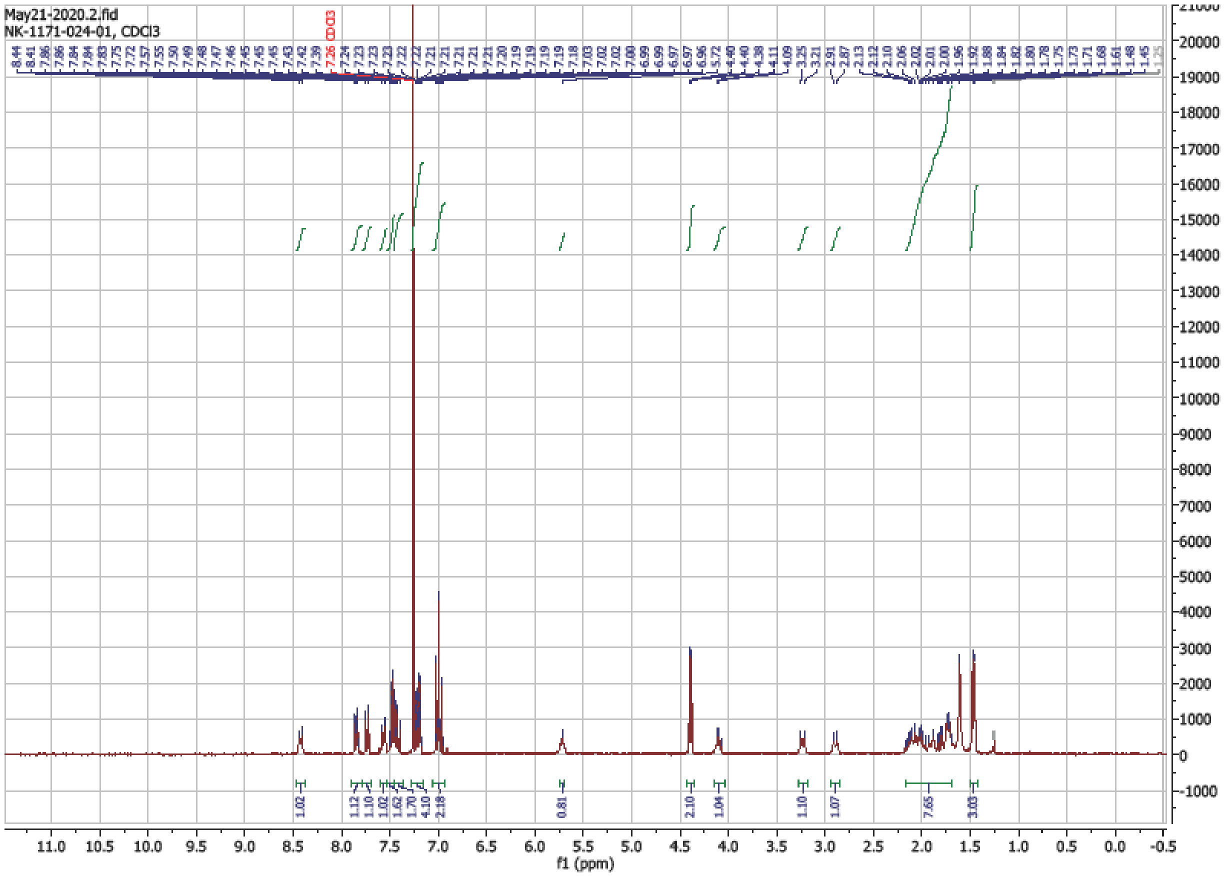

#### Rac3k

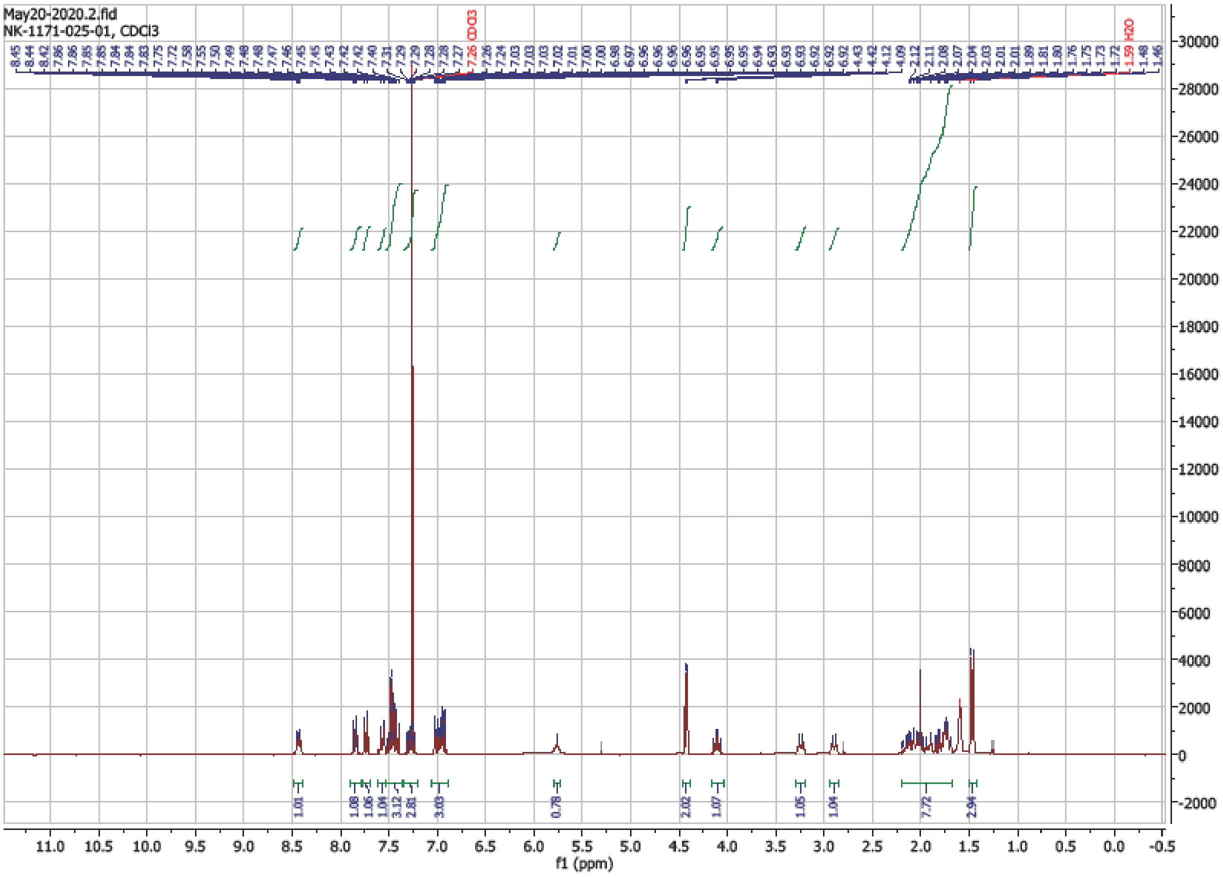

### Rac5c

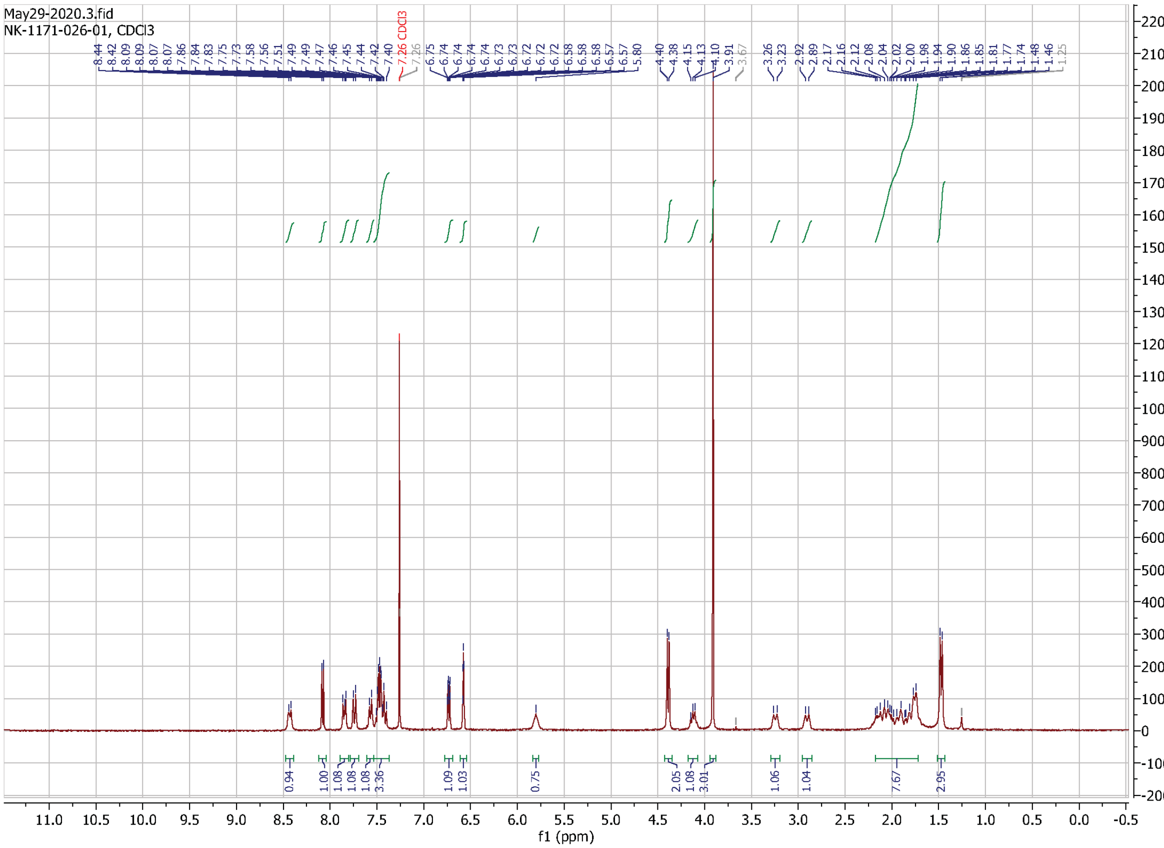
